# Cardiac signals shape insular cortex activity and emotion coding

**DOI:** 10.64898/2026.03.18.712676

**Authors:** Meryl Malezieux, Jeong Yeongseok, Andrea Ressle, Bianca Schmid, Nadine Gogolla

## Abstract

Accumulating evidence suggests that cardiac information influences neuronal activity and fundamental brain functions across species. Yet, how the brain represents cardiovascular signals and how this may impact the processing of emotion states remains unclear. Here, we show that intracellular dynamics and single unit activity in the posterior insular cortex (pInsCtx) are precisely regulated by cardiac signals, with individual neurons tuned to heartbeats in a frequency-dependent manner. Furthermore, heartbeat tuning in the pInsCtx occurred preferentially during the first phase of the cardiac cycle, at systole. Heartbeat tuning increased during both appetitive and aversive emotion states, a process not simply explained by increases in heart rate. Manipulation of sympathetic cardiac arousal using beta-adrenergic blockade disrupted neuronal encoding of emotion states in the pInsCtx and blunted behavioral and bodily emotion expression. These findings reveal precise sensory coding of cardiovascular signals in the pInsCtx, which supports its role in encoding emotion states.

## Introduction

A growing body of evidence demonstrates that sensory information from the heart can affect fundamental brain functions across species. This is indicated by direct effects of heart rate changes on emotion, such as anxiety behavior or fear extinction learning in rodents.^1,2^ Other findings have highlighted that changes in heart rate influence sleep,^3^ and that the timing of different phases of the cardiac cycle shape cognitive functions.^4–6^

However, our understanding of how sensory information from the heart may influence brain function is limited. Importantly, we do not know how the brain represents cardiovascular signals and how this information influences other mental processes such as emotions.

A critical hub for cardioception and emotion is the insular cortex. Indeed, the insular cortex receives ascending sensory information from the heart via polysynaptic afferents,^7^ but also influences heart rate and blood pressure via top-down circuits.^8,9^ Additionally, recent studies in mice have highlighted causal links between cardioception and the regulation of emotion states in the posterior (‘visceral’) insular cortex (pInsCtx).^1,2^ However, how sensory information from the heart is represented in the pInsCtx and how this affects its functions, such as emotion processing, has not been investigated.

Here, we sought to assess how cardiac signals are encoded in the mouse pInsCtx under basal conditions and during emotion states. We first probed the precise neural representation of cardiac variables in the pInsCtx using intracellular and silicon probe electrophysiological recordings paired with readouts of cardiac function, as well as arousal and locomotion. We then explored the impact of emotion on cardioception in the pInsCtx, before assessing how disrupting cardiac signals affects one of the core functions of the pInsCtx: its capacity to encode emotion states.

## Results

### pInsCtx membrane potential dynamics reflect heart rate changes

Previous studies found that strong changes in heart rate during fear-like states influence neuronal activity in the InsCtx of mice^1,2^ and humans.^10^ However, so far, no study has characterized how heart rate is reflected in InsCtx single neuron activity. To address this, we first wondered if cardiac signals may directly shape intracellular dynamics of pInsCtx neurons. We performed whole-cell patch clamp recordings of pInsCtx neurons together with ECG, pupil, and locomotion measurements in head-fixed mice (**Fig. 1A-C**). Interestingly, pInsCtx neurons consistently depolarized when heart rate increased (**Fig. 1D**) and pInsCtx membrane potential correlation with heart rate was stronger than with pupil dynamics or locomotion (**Fig. 1E**), suggesting a specific coupling between cardiac signals and subthreshold activity of insular neurons.

### pInsCtx membrane potential oscillates with the cardiac cycle

Previous work shows that individual heartbeats can affect neuronal activity,^11,12^ therefore suggesting that the timing of heartbeats may impact membrane potential dynamics as well. To address this, we aligned membrane potential fluctuations to the occurrence of heartbeats in the ECG signal (**Fig 1F-G**). This revealed that the membrane potential of pInsCtx neurons oscillated closely together with the cardiac cycle (**Fig. 1G, H**). Cross-correlation indicated that the cardiac oscillation slightly preceded the membrane potential oscillation by a negative lag of -4.2ms (**Fig. 1G, bottom**), suggesting a bottom-up influence of the cardiac cycle on the membrane potential of pInsCtx neurons. Power spectrum analysis revealed a peak frequency around 10 Hz for both the cardiac cycle and membrane potential oscillations (**Fig. 1H**). To control for the specificity of the membrane potential oscillation to the cardiac cycle, we shuffled the timing of the detected heartbeats by randomizing the detection within the recording. This resulted in a loss of intracellular oscillations, showing that this oscillation is indeed synchronized with the cardiac cycle and is only apparent when aligned to the true heartbeats (**Extended Data Fig. 1**).

**Figure 1.**
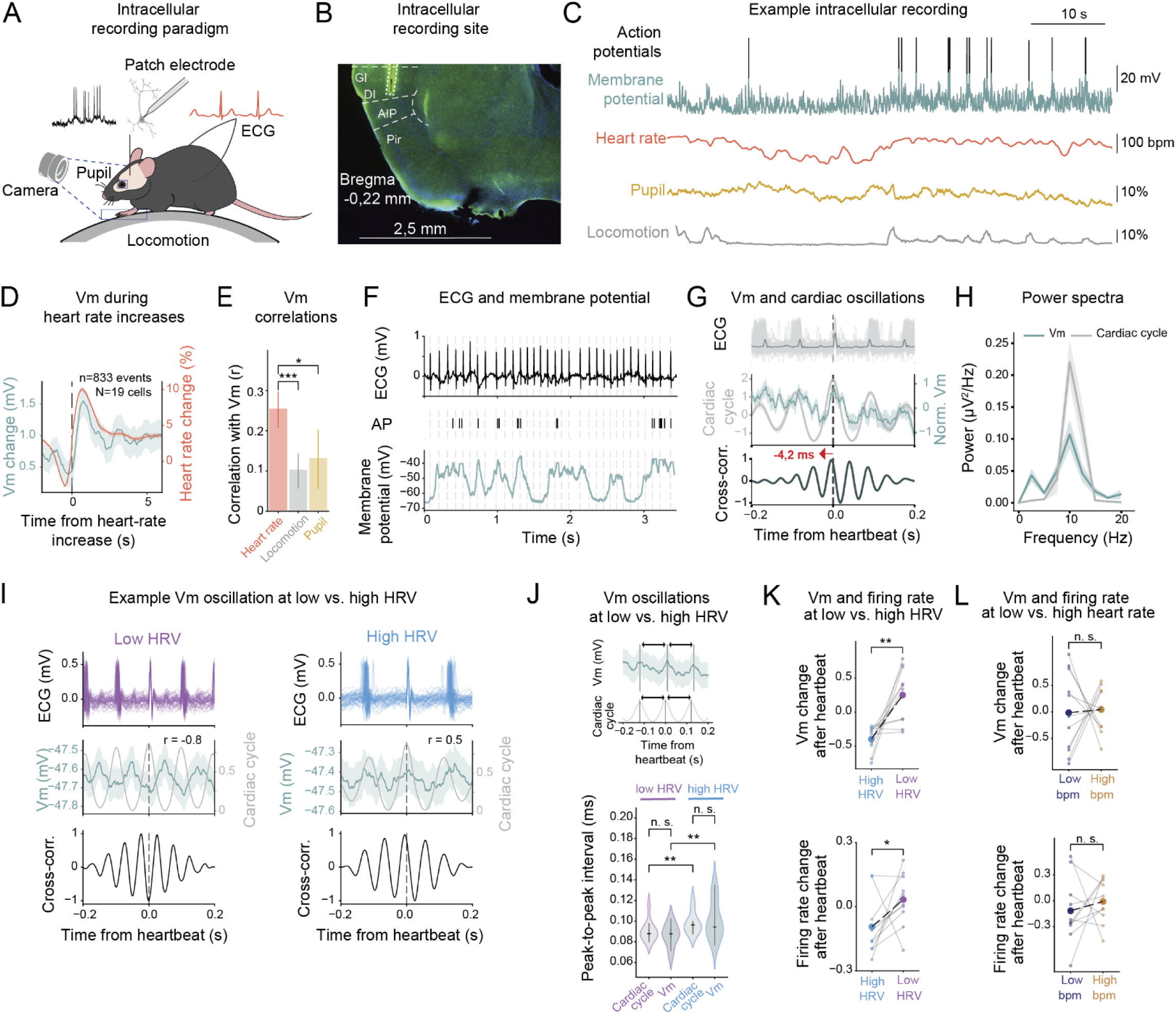
Cardiac signals modulate the membrane potential of pInsCtx neurons. **A.** Schematic of whole-cell patch clamp recordings in the pInsCtx of head-fixed mice with tethered ECG monitoring and video recording of locomotory behavior and pupil dynamics. **B.** Epifluorescence microscope image of a coronal section of pInsCtx, showing the track of the patch electrode in green and delineated by a dotted rectangle. **C.** Example patch clamp recording of a pInsCtx neuron showing the membrane potential (Vm) fluctuations below (top, green) and above (top, black) action potential firing, with heart rate (red), pupil size (yellow) and locomotion (grey). **D.** Overlay of heart rate increases (red) with the corresponding depolarization in membrane potential (green) relative to increases in heart rate (0s). Traces represent mean ± s.e.m. **E.** Pearson’s correlation coefficients of membrane potential to heart rate, locomotion or pupil size. (Friedman test (p<0.05) followed by post hoc Wilcoxon test, *p < 0.05, ***p<0.001). Data is represented by median ± s.e.m. for the positive correlation coefficients (n=17/19 cells in N=12 mice). **F.** Example of an ECG recording (top) and the corresponding action potentials (middle), and membrane potential (bottom) of an example cell. Every dashed line shows a detected heartbeat. **G.** pInsCtx membrane potential oscillates relative to heartbeats. Top: ECG centered around each heartbeat at 0 s overlaid with the average ECG. Middle, corresponding cardiac cycle (grey, peaks represent R peaks times) overlaid with the membrane potential fluctuations (green) relative to heartbeats. Traces are average signals ± s.e.m. of 10 cells in 7 mice. Bottom: cross-correlation of the cardiac cycle with the membrane potential. The maximum cross-correlation (0.95) occurs with a negative lag of -4.2ms. **H.** Power spectral density of the cardiac cycle (grey) and the membrane potential (green) around heartbeats. The cardiac cycle power peaks at 10Hz, similar to the membrane potential. Traces represent average ± s.e.m. of 10 cells in 7 mice. **I.** Membrane potential relative to the cardiac cycle for low HRV (left, purple) and high HRV (right, blue). Top: ECG centered around each heartbeat at 0 s. Middle: corresponding cardiac cycle (grey, peaks represent R peaks times) overlaid with the membrane potential fluctuations (green) relative to heartbeats. Traces are average signals ± s.e.m. of 10 cells in 7 mice. Bottom: cross-correlation of the cardiac cycle with the membrane potential. Middle, left: when HRV is low, the membrane potential and cardiac cycle are anticorrelated (r = -0.8). Middle, right: when HRV is high, the membrane potential and cardiac cycle are positively correlated (r = 0.5). **J.** Top: example membrane potential (top) and cardiac cycle (bottom). Arrows indicate peak-to-peak intervals, extracted measures compared on the bottom plot. Bottom: distributions of the peak-to-peak intervals in ms during low HRV (purple) and high HRV (blue) for the cardiac cycles and membrane potential oscillations (Vm). Cardiac cycles and membrane potential oscillations peak-to-peak intervals were shorter during low HRV as compared to high HRV (Friedman test (p<0.05) followed by post hoc Wilcoxon test, **p < 0.005). No difference between the peak-to-peak intervals duration were found between cardiac cycle and membrane potential oscillations during low HRV, nor during high HRV. **K.** Membrane potential changes (top) and firing rate changes (bottom) relative to heartbeats during high vs. low HRV for 10 cells in 7 mice (student t-test, top: **p<0.01, bottom: *p<0.05). **L.** Membrane potential changes (top) and firing rate changes (bottom) relative to heartbeats during high vs. low heart rate for 10 cells in 7 mice (student t-test, n.s.).

### Cardiac influence on pInsCtx membrane potential is state-dependent

Cardiac activity can be assessed through heart rate or heart rate variability (HRV). While heart rate refers to the number of heartbeats per minute, HRV corresponds to the variations in the time intervals between consecutive heartbeats. Interestingly, HRV is known to reflect autonomic balance and decreases in states of stress or strong emotion.^13^ We argued that if cardiac sensory signals truly influence the membrane potential of pInsCtx neurons, then the duration of the oscillations in membrane potential should follow HRV fluctuations. Indeed, our analysis revealed that when the cardiac cycle was either long or short, the membrane potential oscillations lengthened or shortened, respectively (**Fig. 1I-J**). Interestingly, we observed an unexpected phase shift of membrane potential oscillations around the cardiac cycle (**Fig. 1I**). When HRV was low, the membrane potential oscillation was anti-correlated with the cardiac cycle, and when HRV was high, the membrane potential oscillation was positively correlated with the cardiac cycle (**Fig. 1I**). This translated to a depolarization and an increase in firing rate after heartbeats when HRV was low, and a hyperpolarization and a decrease in firing rate after heartbeats when HRV was high (**Fig. 1K**). Interestingly, this modulation was not evident during periods of low and high heart rates (**Fig. 1L**).

### pInsCtx neurons are tuned to heart rate and individual heartbeats

Our findings demonstrate that the membrane potential of individual cells in the pInsCtx precisely fluctuates with the cardiac cycle in a state-dependent manner. Based on these observations, we wondered whether this cardiac modulation could also impact population activity in the pInsCtx.

Using silicon probe recordings (**Fig. 2A-B, Extended Data Fig. 2A**), we first observed that, similar to their membrane potential, pInsCtx neurons consistently increased firing upon increases in heart rate. This modulation was not observed in neurons in the whisker motor cortex (wM1), suggesting that this property is not shared between all cortical regions (**Extended Data Fig. 2B, Extended Data Fig. 3**). Importantly, 22.9% of insula neurons were still modulated by heart rate increases in the absence of locomotion, and the average amplitude of the increase in activity was similar during immobility and locomotion, suggesting that heart rate can influence neuronal activity in the pInsCtx independently of locomotion (**Extended Data Fig. 2C**). Using a L1-linear regularized Lasso regression, we found that heart rate was predicted with significantly higher accuracy (0.66 ± 0.08) from pInsCtx activity than locomotion (0.41 ± 0.09) or pupil (0.43 ± 0.07) (**Extended Data Fig. 2D**), indicating that among these variables, heart rate is the predominant factor shaping population activity in the pInsCtx. Finally, we quantified the proportion of neurons selective to increases in heart rate, pupil, or locomotion. Again, the largest fraction of neurons with unique selectivity was responsive to heart rate only (**Extended Data Fig. 2E**).

Having observed a strong modulation by heart rate and a precise membrane potential oscillation with the cardiac cycle, we next asked whether pInsCtx neurons could also be modulated by individual heartbeats.^11^ We detected each action potential relative to the closest occurring heartbeat (**Fig. 2C**) and computed the phase of action potentials relative to heartbeats. We then performed a Rayleigh circular test on the corresponding phases. This test identifies if a population is distributed uniformly around a circle.^14^ A low p-value (< 0.05) suggests that the unit fires at a consistent time interval after the heart beats. We found that a proportion of pInsCtx units (58/499 neurons) displayed stereotyped firing rate around heartbeats (**Fig. 2D, E**). Across recordings, 11.1% of the total number of pInsCtx neurons were tuned to heartbeats (**Fig. 2E**).

**Figure 2.**
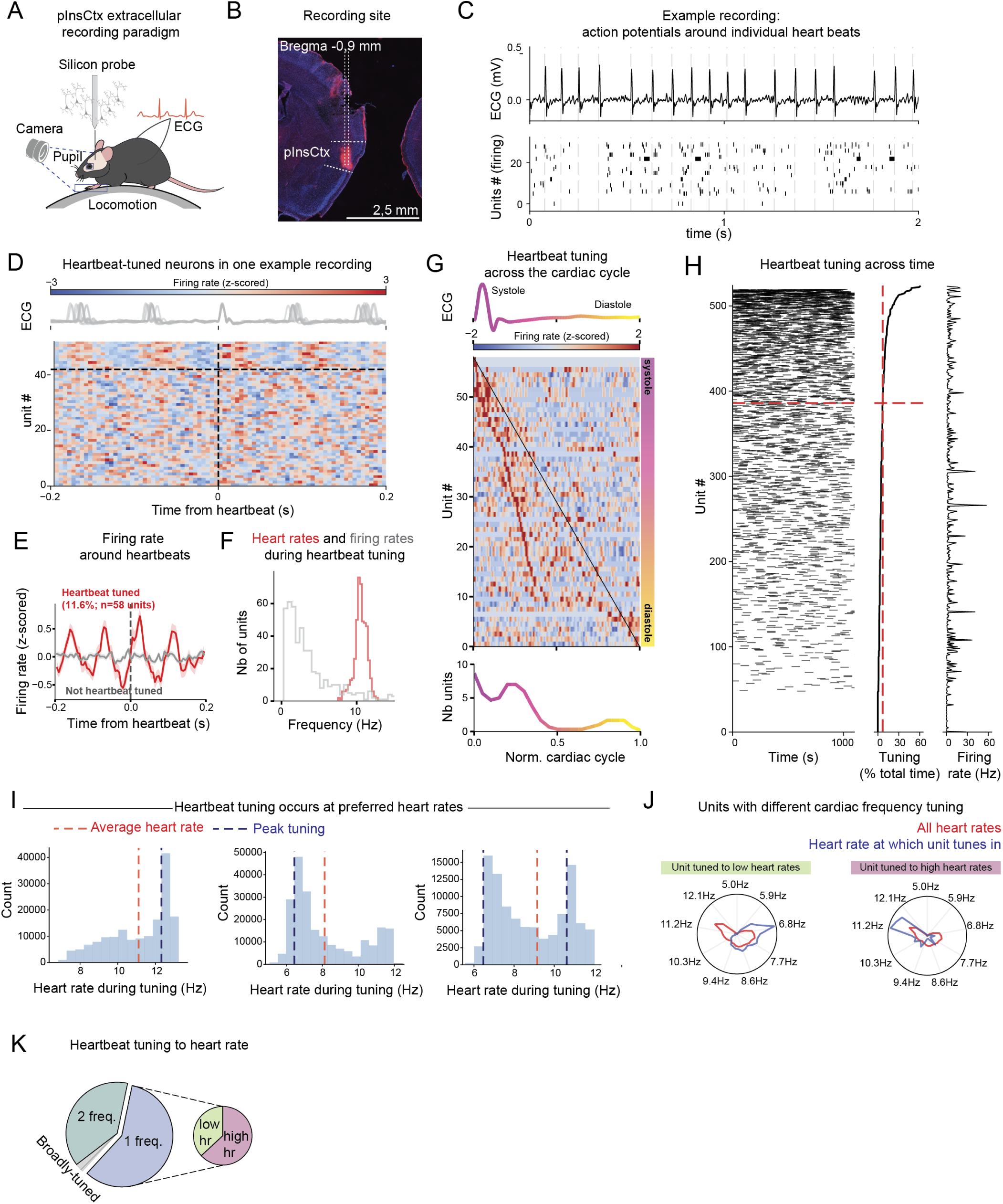
pInsCtx neurons are tuned to single heartbeats. **A.** Schematic of simultaneous silicon probe recordings in the pInsCtx, tethered ECG monitoring, and video recording of locomotion and pupil dynamics in head-fixed mice. **B.** Epifluorescence microscope image showing the silicon probes track in pink (labelled with DiI and approximate probe track as dotted line) on a coronal brain section at the level of pInsCtx. **C.** Detection of action potentials around heartbeats. Top: ECG Bottom: raster of action potentials of an example recording. Each dashed line represents a detected heartbeat. **D.** Example recording showing 19.2% (10/52) of neurons significantly tuned to heartbeats (above the dashed line) (rayleigh test, *p<0.05). Heatmap represents z-scored firing rate of each unit relative to heartbeats during the recording. ECG is shown at the top and units are sorted by significance of heartbeat tuning. **E.** Averaged z-scored firing rate relative to heartbeats for all significantly heartbeat tuned neurons (rayleigh test, *p<0.05; 11.6%, n=58 units in N=14 recording sessions in 10 mice) compared to untuned neurons (88.4%, n=441 units in N=14 recording sessions in 10 mice). **F.** Histogram distribution of firing rates (grey) and heartrates (red) during heartbeat tuning. **G.** Heatmap of all significantly heartbeat tuned neurons sorted according to the normalized cardiac cycle. (top) color-coded ECG with the R peak starting at 0 and the next R peak occurring at 1. (middle) heatmap of the firing rate of all significantly heartbeat tuned neurons sorted according to their peak firing rate during the cardiac cycle (bottom) counts of heartbeat tuned units across the cardiac cycle, color-coded from dark pink (beginning of R-peak) to yellow (end of the cardiac cycle). **H.** (left) raster plot of all recorded neurons (n=524 in N=10 mice) showing heartbeat tuning events throughout the recordings. Each black line represents a neuron getting tuned to the heartbeat during a 10s timebin. The neurons are sorted according to the duration spent tuned, with the highly tuned neurons at the top and the low tuned neurons at the bottom. (middle) corresponding time spent heartbeat tuned during the recording (right) corresponding firing rate for each neuron. The red dashed lines correspond to the average time spent heartbeat tuned (6.9% total recording time). **I.** Examples distributions of heart rates during heartbeat tuning for 3 different units. The red dashed line corresponds to the average heart rate during the recording. The blue dashed line corresponds to the peak in heartbeat tuning. **J.** Polar plots of example units with distinct tuning properties for cardiac frequencies. (left) example unit tuned to low heartrates (blue) compared to the distribution of all heartrates (red) (right) same as (left) but for a unit tuned to high heartrates. **K.** Pie-chart showing the proportion of neurons having a single heart frequency preference for tuning (58.6%), or two frequencies (38.7%), or more than two frequencies (2.7%). Among the neurons that have a single preferred frequency, 63.1% tune in when heartrate is high, while 36.9% tune in when heartrate is low.

To provide further evidence that heartbeat tuning does not occur due to stochastic coincidence between neuronal firing rate and heart rate, we computed the frequencies of heart rate and neuronal firing during heartbeat tuning. We found that heart rates and firing rates during heartbeat tuning did not overlap, suggesting that neurons are not coincidentally tuning to heartbeats because they are firing at the same frequency as the heart is beating (**Fig. 2F**).

To investigate if heartbeat tuning was specific to pInsCtx neurons, we performed the same analysis for single units recorded in wM1. We observed that only 4.7% of wM1 units were tuned to heartbeats, similar to randomly generated spike trains (4.4%) (**Extended Data Fig. 4**).

### Heartbeat tuning occurs preferentially at systole

Among the neurons that were tuned to heartbeats, we noticed that different neurons were active at staggered latencies after heartbeats (**Fig. 2D**). Thus, we asked whether individual neurons were tuned to specific timepoints within the cardiac cycle. We normalized the cardiac cycle durations: each cardiac cycle was rescaled from 0 to 1 (arbitrary units), with 0 corresponding to the onset of the cycle (e.g., R-peak in ECG or systolic onset), and 1 corresponding to the completion of the cycle (onset of the subsequent R-peak). We then sorted the heartbeat tuned neurons by their peak firing rate within the cardiac cycle, which is comprised of two main phases: systole and diastole (**Fig. 2G**). This sorting highlighted that the majority (74%) of the heartbeat tuned units fired during systole, while the remaining 26% spanned the other two-thirds of the cardiac cycle. Taken together, these results suggest that neurons in the pInsCtx are strongly entrained when the heart is beating, with a denser coding during the contraction of the heart chambers and increase in arterial blood pressure.

### A majority of pInsCtx neurons can be heartbeat tuned

We next wondered whether heartbeat tuning was a stable coding property of the identified subset of 10-12% of insular neurons. However, we found that across an entire recording session of approx. 20 min., individual insular neurons did not exhibit stable heartbeat tuning but rather intermittently tuned in and out of the heartbeat (**Fig. 2H, left**). Individual neurons exhibited large differences in the overall time they spent heartbeat tuned, varying between 0 up to 60% (6.9% ±0.4% of the total recording time on average, **Fig. 2H, center; Extended Data Fig. 5A**). Most strikingly, the great majority of recorded neurons, up to 85% percent (**Extended Data Fig. 5B**), exhibited at least one 10 s epoch of heartbeat tuning during the recording session, suggesting that heartbeat tuning is a common property of most neurons in the pInsCtx. Importantly, single units that exhibited frequent versus infrequent heartbeat tuning did not differ substantially in firing rates, indicating that frequent heartbeat tuning was not due to high firing rates (**Fig. 2H, right**). To control that heartbeat tuning is not sensitive to the chosen timebin, we computed heartbeat tuning across time for each pInsCtx unit for several bin sizes. Interestingly, we found that the different bin sizes, ranging from 200ms to 10s, did not significantly affect heartbeat tuning, with the general structure of tuning per unit being conserved across bin sizes (**Extended Data Fig. 5C**).

### pInsCtx neurons exhibit precise heartbeat frequency tuning

Because heart rate fluctuates over time and influences many insular neurons, we next asked whether pInsCtx neurons might be tuning in and out at specific cardiac frequencies. Indeed, frequency tuning is a common property of neurons in sensory cortices, including spatial frequency tuning in visual cortex or sound frequency tuning in auditory cortex.^15,16^ To ask whether insular neurons may be tuned to specific heartbeat frequencies, we computed their tuning preference by extracting all heart rates at which each unit tuned in, and then detecting the peak tuning frequency for each neuron (see **Fig. 2I-J** for single unit examples). This revealed that a majority of insular neurons exhibited strong frequency tuning with only 1 preferred (58.6%) or 2 preferred (38.7%) heartbeat frequencies (**Fig. 2K**). Amongst the units which were tuned to a single frequency, a larger proportion (61.4%) tuned in when heartrate was high (above average heart rate) (**Fig. 2K**). Overall, very few neurons were broadly tuned, i.e. tuning into two or more frequencies (**Fig. 2K**).

### Heartbeat tuning requires synaptic transmission

What could be the source of this heartbeat tuning? One possibility may be a recently described local, brain-intrinsic activation of mechanoreceptors which would react to changes in blood pressure from nearby blood vessels.^11^ Alternatively, cardiac signals may be relayed to the pInsCtx via a polysynaptic route, potentially through the vagus nerve, which is known to carry sensory information from many peripheral organs to the brainstem.^7^

To test whether synaptic transmission is necessary for heartbeat tuning in the pInsCtx, we locally infused antagonists of AMPA and NMDA glutamatergic receptors, as well as nicotinic acetylcholine receptors (CNQX 1mM, APV 2mM and Mecamylamine 1mM) next to the silicon probes (**Extended Data Fig. 6A**). We argued that if heartbeat tuning occurs solely via mechanoreceptors, and not via synaptic transmission, we should still observe heartbeat-evoked neuronal firing in neurons whose activity decreases due to the block of synaptic input. However, while we observed that, as expected, blocking glutamatergic and cholinergic transmission resulted in marked reduction in the activity of individual pInsCtx neurons, heartbeat tuning also decreased proportionally in the affected neurons. Only in the non-affected, active population, the proportion of heartbeat tuned neurons remained stable (11.1% ±3.0 before blockade to 14.2% ±7.6 after blockade heartbeat tuned units; **Extended Data Fig. 6B-D**). While not fully excluding the involvement of mechanoreceptors, these results suggest that heartbeat tuning in the insula is at least partially dependent on synaptic transmission.

### Heartbeat tuning increases during emotion states

What may be the functional implications of the cardiac modulation of pInsCtx activity? Given the role of the insular cortex in interoception and emotion regulation,^1,17–19^ we wondered how the cardiac modulation of pInsCtx activity described here may influence emotion processing. Recent evidence suggests that experimentally-induced tachycardia can elicit anxiety and is dampened by pInsCtx inhibition.^1^ However, how physiological reactivity of the heart to emotion may be represented in the pInsCtx and influence its capacity to process emotion remains unclear.

**Figure 3.**
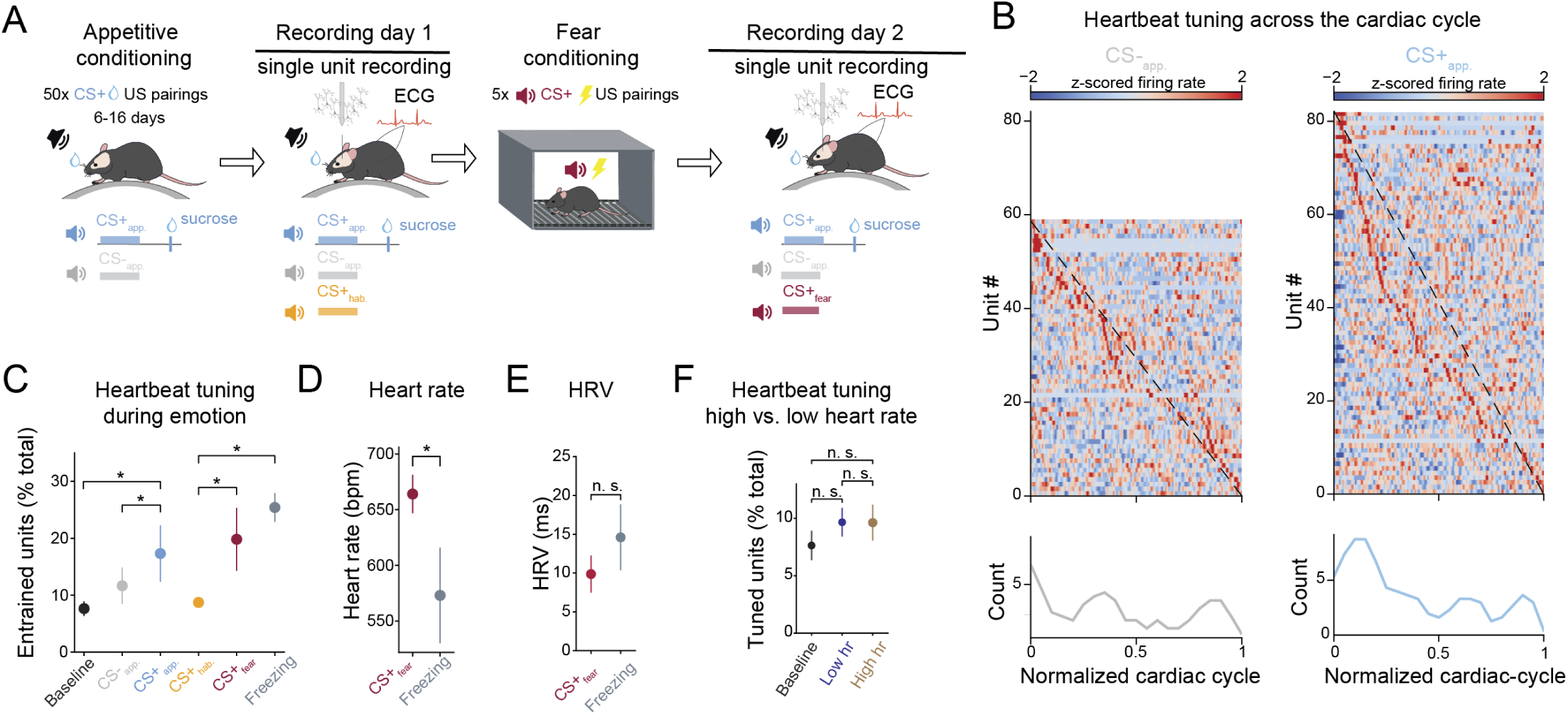
Heartbeat tuning of pInsCtx neurons is enhanced during emotion states. **A.** Schematic of the experimental timeline. Left: Mice were first trained in an appetitive conditioning paradigm to associate a CS+_app_ with the delivery of a reward (sucrose), and to discriminate between the CS+_app_ and CS-_app_. Appetitive training occurred once a day for 6-16 days. Once successfully trained, mice entered the first recording day and exposed to CS+_app_ and CS-_app_. They were further exposed to a third, novel CS+ (CS+_hab_). On that same day and after the recording session, mice were placed in a fear conditioning chamber where the CS+_hab_ was paired with a foot shock (US) 5 times. After fear conditioning, the CS+_hab_ was referred to as CS+_fear_. On the next day, mice entered the second recording day and exposed to CS+_app_, CS-_app_, as well as CS+_fear_. **B.** Heatmap of the z-scored firing rate of all units that are significantly heartbeat tuned (*p<0.05, rayleigh test) during CS-_app_ (left) and appetitive CS+ _app_ (right). Units are sorted according to their peak firing rate in the normalized cardiac cycle. **C.** Quantification of the percentage of heartbeat tuned units during baseline, CS-_app._, CS+ _app.,_ CS+_hab._, CS+_fear_, and freezing. (Wilcoxon and Mann-Whitney U-tests, *p<0.05). Data represents mean ± s.e.m. **D.** Average heart rate during CS+_fear_ and freezing (student t-test, *p<0.05). Data represents mean variable ± s.e.m. N=4 mice. **E.** Average HRV during CS+_fear_ and freezing (student t-test, n.s.). Data represents mean variable ± s.e.m. N=4 mice. **F.** Percentage of heartbeat tuned pInsCtx units during baseline (black), periods of low heart rate (blue), periods of high heart rate (brown). Baseline periods were defined as three 15s epochs before any CS presentation. Low and high heart rate epochs were defined as three 15s continuous periods were heart rate was below or above the average heart rate respectively, and outside of any CS presentation, freezing event, or baseline epoch. Data represents mean variable ± s.e.m.

To trigger emotion states of opposite valence, we performed appetitive and aversive conditioning within the same animal (**Fig. 3A**). Head-fixed mice first underwent appetitive conditioning (AC), where they learned to associate an auditory cue (CS+_app_) with the delivery of sucrose (10% diluted in water), while a second tone, CS-_app_, was not followed by reward. Successful learning was reflected by increased licking rates in response to the CS+_app_ (**Extended Data Fig. 7A**). After successful AC acquisition, we performed a first recording session, during which mice were also habituated to a third tone (CS_hab_). Following the first recording session, mice underwent a classical fear conditioning (FC) paradigm in a separate chamber to associate the previous CS+_hab_, now CS+_fear_, to the delivery of a mild footshock. During FC, mice displayed increased freezing behavior upon pairing CS+_fear_ with footshock presentations (**Extended Data Fig. 7B**). A day after FC, we performed a second, identical head-fixed recording session (**Fig. 3A**).

Emotions have been found to induce heart rate increases and pupil dilations as signs of heightened emotional arousal in human and nonhuman animals.^20,21^ Furthermore, pInsCtx activity has been shown to respond to emotionally valenced cues.^18,22^ Accordingly, in our paradigm, CS+ presentations of both valences triggered increases in heart rate, pupil diameter, locomotion and pInsCtx activity, suggesting the successful induction of emotional states through the presentation of the associative cues (**Extended Data Fig. 7C-D**). In addition to the emotional states induced via the presentation of associative cues, we also assessed whether freezing, as behavioral indicator of a fearful state, induced consistent bodily, behavioral and insula activity changes. In agreement with previous studies,^2,23^ freezing episodes were associated with decreased pInsCtx firing and heart rate, and increased pupil size in the absence of locomotion (**Extended Data Fig. 7E**).

Having established a paradigm that induced emotional arousal during CS+ presentations of different valences and during freezing, we next sought to determine how emotional states may affect heartbeat tuning in the pInsCtx.

We first assessed how many neurons in the pInsCtx were heartbeat tuned during emotion states and found that the percentage of heartbeat tuned neurons was higher during emotion, namely during fearful freezing, and aversive and appetitive CS+ presentations, compared to the control cues CS-_app_ or CS_hab_ (**Fig. 3B-C**). The fact that most heartbeat tuned neurons were observed during fearful freezing, when the heart rate consistently decreased and was lower than during the CS+_fear_ (**Fig. 3D-E**) indicated that the increase in heartbeat tuned neurons was not simply a consequence of higher heart rates during emotional arousal. To test this further, we compared the amount of heartbeat tuned neurons during baseline, versus low and high heart rates. We separated our data into baseline epochs, before any emotional cue was presented, and high versus low heart rates epochs during the emotion paradigm but outside of emotional cues or freezing events. We found no significant differences in the percentage of heartbeat tuned neurons (**Fig. 3F**), further confirming that heartbeat tuning during emotion states does not simply increase due to higher heart rates.

Together, these results demonstrate that heartbeat tuning in pInsCtx neurons is specifically enhanced during emotion states (**Fig. 3B**).

### Systemic blockade of beta 1 adrenergic transmission decouples pInsCtx firing from heart rate

Emotional arousal leads to sympathetic activation and release of adrenaline and noradrenaline in the body, which via the binding to ß1-adrenergic receptors in the heart increases heart rate and the strength of cardiac contractions.^24^ We hypothesized that the sudden, conditioned increase in rate and strength of cardiac contractions induced through sympathetic activation may lead to the increase in heartbeat tuning in the insula during emotion. To test this hypothesis directly, we utilized a beta-blocker, metoprolol, which primarily acts by blocking ß1-adrenergic receptors in the heart. Accordingly, upon systemic metoprolol administration, we observed a slight but long-lasting reduction in heart rate (**Fig. 4A-B**). However, we observed no significant changes in pInsCtx firing rate (**Fig. 4C**), HRV, locomotion, or pupil size (**Extended Data Fig. 8A-C**). Importantly, under baseline conditions and without explicit emotion induction, heartbeat tuning in the pInsCtx was unaffected by metoprolol (**Extended Data Fig. 8D**).

We here described that heart rate increases were correlated to increases in firing rates in the pInsCtx under baseline conditions (**Extended Data Fig. 2B-C**). Strikingly, we found that under metoprolol, this correlation was absent (**Fig. 4D**). Interestingly, this decorrelation was accompanied by a decreased predictive power of heart rate from pInsCtx activity, while the decoding accuracies for pupil and locomotion remained unchanged (**Extended Data Fig. 9**), suggesting a specific loss of cardiac signal representation in pInsCtx. While we observed an overall shift towards smaller heart rate increases (**Extended Data Fig. 8E**), overall heart rates distribution differed mostly in the absence of high heart rates under metoprolol (**Extended Data Fig. 8F**). Reduced increases in heart rate led to much smaller increases in firing rate, although general firing rate of pInsCtx was not affected by metoprolol (**Extended Data Fig. 8G-H**).

### Systemic blockade of beta 1 adrenergic transmission disrupts emotion processing in the insular cortex

Given its effects on adrenergic transmission, we hypothesized that metoprolol would lead to reduced emotional arousal in response to valenced cues. Indeed, while in control mice heart rate strongly increased upon CS+ presentations (CS+_app_ and CS+_fear_), these increases were much weaker under metoprolol (**Fig. 4E-F; Extended Data Fig. 10A**), demonstrating a dampening effect of sympathetic arousal on the heart via the beta blocker.

InsCtx neurons respond strongly to learned associative cues (**Fig. 4G-H; S7C-D**).^2,18,22,25^ Strikingly, these neuronal responses were abolished under metoprolol (**Fig. 4G-H**). However, the lack of response in firing rate was limited to the associative cue presentations, leaving intact both the US response to sucrose (**Extended Data Fig. 10C**) and the sensory responses to individual tone pips (**Extended Data Fig. 10D**) comprising the auditory CS+ _fear_. This suggests that, while metoprolol did not affect neuronal firing in general (**Fig. 4C**), or sensory evoked activity, it had strong selective effects on abolishing pInsCtx activity in response to emotionally valenced cues. Additionally, metoprolol affected the behavioral and physiological responses to valenced cues (CS+_app_ and CS+_fear)_, but not to CS-, assessed by changes in pupil size, locomotion as well as licking for the CS+_app_ (**Extended Data Fig. 10E-G**).

These results demonstrate that preventing heart rate changes via beta blockade impairs pInsCtx neuronal responses as well as physiological and behavioral responses to emotional cues.

### Metoprolol decreases heartbeat tuning during emotion states

Given that metoprolol prevented the insula to respond to associative cues, we hypothesized that this may be due to the decoupling of heart rate and pInsCtx activity under metoprolol (**Fig. 4D**). Thus, we next analyzed whether heartbeat tuning in pInsCtx neurons was affected under metoprolol. We found that during CS+_app_ and CS+_fear_ presentations, the percentage of heartbeat tuned neurons in the pInsCtx was significantly reduced (**Fig. 4I, Extended Data Fig. 10H, J**). However, no difference in heartbeat tuning was observed for the CS-_app_ (**Fig. 4I, Extended Data Fig. 10I**), suggesting that beta blockade specifically disrupts heartbeat tuning during emotional cues.

### Metoprolol impairs emotion state coding in pInsCtx

pInsCtx units did not tune to the heartbeat and did not respond to emotional cues under β-blockade. We therefore investigated how β-blockade affected the capacity of pInsCtx to decode specific emotional cues and states. We performed PCA on the activity of all units during valenced cues and behaviors. We used the activities upon CS+ and CS-presentations, as well as pInsCtx activity during licking and freezing, as examples of appetitive and aversive behavioral expressions, respectively.

We observed that, after fear conditioning in control animals, confidence interval ellipses were clustered separately along the first two principal components (PCs) (**Fig. 4J**). However, under metoprolol, the ellipses collapsed into the same PC space and overlapped (**Fig. 4K**). Additionally, the variance explained by each PC was reduced by a factor of approximately two (**Fig. 4J-K**). Together, this suggests that metoprolol disrupts the neural representations of emotional valence in the pInsCtx.

To corroborate these findings, we trained an SVM linear classifier to decode CS identities from pInsCtx activity (**Fig. 4L-O**). Under control conditions, CS identity (CS+_app_ vs. CS-_app_) was accurately decoded from pInsCtx activity during the CS presentation. However, under metoprolol, CS identity could not be accurately decoded anymore. Notably, after sucrose delivery, the trial type was accurately decoded, suggesting that metoprolol specifically affected the associative affective component triggered by the cue (**Fig. 4L**).

**Figure 4.**
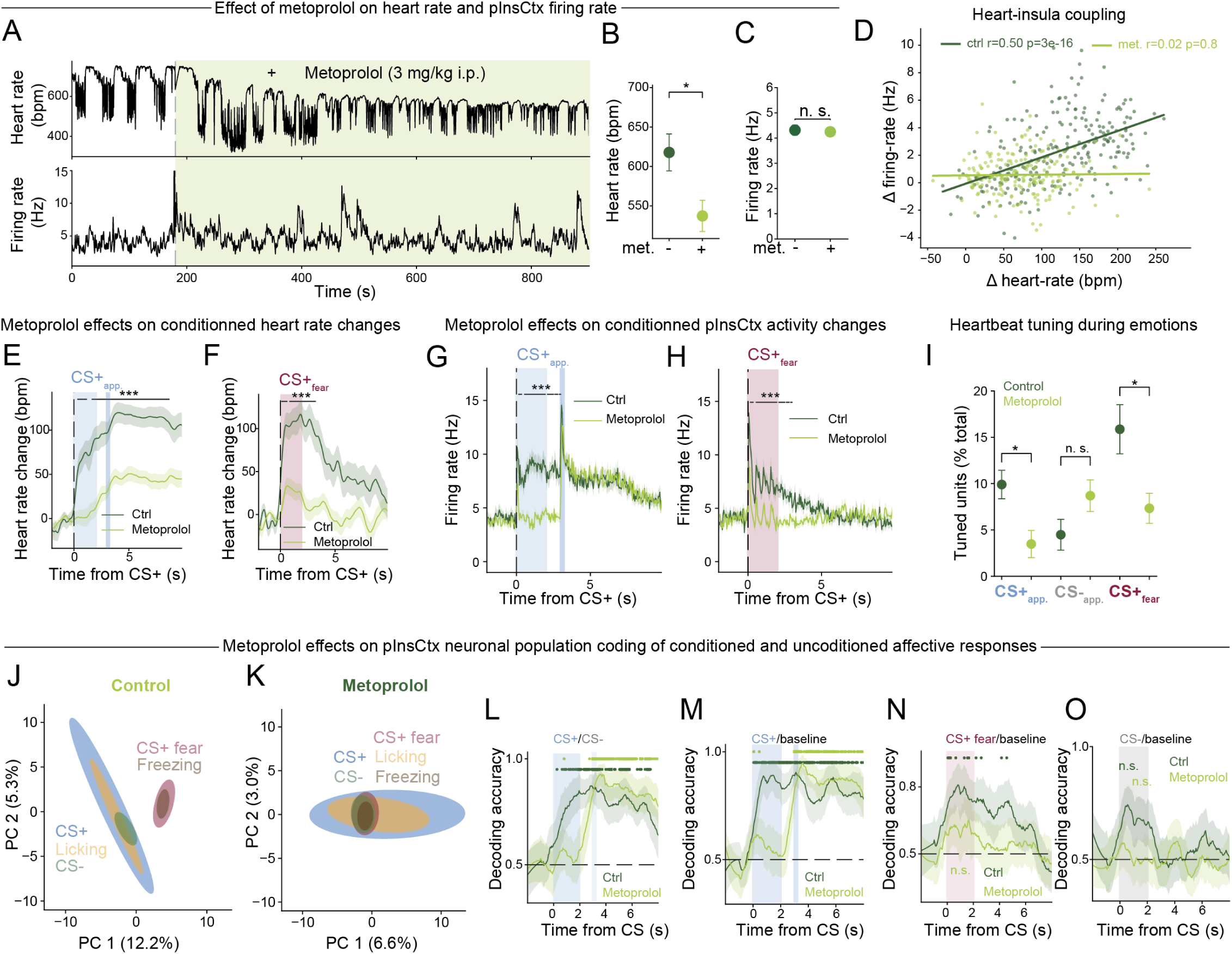
Metoprolol reduces cardio-insular coupling and prevents emotion state coding in the pInsCtx. **A.** Example heart rate (top) and pInsCtx neuronal population average firing rate (bottom) illustrating the effect of metoprolol administered at 180s. **B.** Heart rate before and after metoprolol administration (student t-test, *p<0.05). Data represents mean ± s.e.m. N=4 mice. **C.** pInsCtx single unit average firing rate before and after metoprolol administration (student t-test, n.s.). Data represents mean ± s.e.m. N=4 mice. **D.** Correlation between heart rate increases and the corresponding pInsCtx firing rate increases in control (n=237, day 1, dark green) versus metoprolol-treated (n=201, day 2, light green) mice. Each dot corresponds to an increase in heart rate. N=4 mice per group. **E.** Heart rate change relative to the CS+_app._ in control (day1) versus metoprolol (day2) groups. Traces are normalized to 2s before the start of the CS. (***p<0.001, repeated measures ANOVA with Bonferroni correction). N=4 mice per group. Data represents mean heart rate ± s.e.m. **F.** Same as in (E) but for CS+_fear_. Data represents mean heart rate ± s.e.m. N=4 mice per group. **G.** Firing rate of pInsCtx neurons relative to CS+_app._ in control versus metoprolol groups (***p<0.001, repeated measures ANOVA with Bonferroni correction). N=4 mice per group. Data represents mean firing rate ± s.e.m. **H.** Same as in (G) but for CS+ _fear_. Data represents mean firing rate ± s.e.m. N=4 mice per group. **I.** Percentage of the total number of pInsCtx units that are heartbeat tuned during the different CS in control and metoprolol treated mice (student t-test, *p<0.05). **J.** Representation of the 2 first components after principal component analysis (PCA) of pInsCtx population activity in control mice after appetitive and fear conditioning during CS presentations and behavioral events. Ellipses represent covariance confidence intervals. **K.** Same as in (J) but for metoprolol treated mice. **L.** SVM classifier decoding accuracies for CS identity (CS+_app_. vs CS-_app_.) from pInsCtx activity (FDR-corrected permutation tests, p < 0.05). **M.** SVM classifier decoding accuracies for CS+_app_. vs baseline from pInsCtx activity (FDR-corrected permutation tests, p < 0.05). **N.** SVM classifier decoding accuracies for CS+_fear_. vs baseline from pInsCtx activity (FDR-corrected permutation tests, p < 0.05). **O.** SVM classifier decoding accuracies for CS-_app_. vs baseline from pInsCtx activity (FDR-corrected permutation tests, n.s.).

Additionally, the CS+_app_ and CS+_fear_ were both accurately decoded from pInsCtx baseline activity in the control but not in the metoprolol group (**Fig. 4M, N**). In contrast, CS-_app_ was never accurately decoded from pInsCtx activity (**Fig. 4O**), suggesting that pInsCtx neurons do not distinctly represent neutral stimuli. These findings demonstrate that pInsCtx neurons encode and differentiate between appetitive and aversive cues, as shown by distinct PCA clusters and accurate SVM decoding of CS+ identities. However, metoprolol disrupts this coding and impairs real-time discrimination of emotional cues.

Thus, blocking beta adrenergic transmission leads to strong deficits in the pInsCtx properties of sensing heart rate changes and individual heartbeats, impairing neural and behavioral representation of emotional states.

## Discussion

The insula serves as a multisensory hub, integrating both exteroceptive and interoceptive signals across timescales. Some of the diverse functions that pInsCtx supports are thermoception, immunoception, interoception, hunger and thirst homeostasis, taste processing, and emotions.^26–32^ Our study broadens our understanding of the interoceptive properties of pInsCtx in the context of emotion state coding.

We found that neurons in the pInsCtx are entrained by individual heartbeats. While the pInsCtx is known as a visceral hub, integrating signals from many peripheral organs,^27,33,34^ how it encodes these signals precisely remained unknown. Our study is the first to (i) reveal precise heartbeat entrainment in pInsCtx neurons, and (ii) uncover a functional role of this cardiac signal processing in emotion.

Our findings highlight that the membrane potential of pInsCtx oscillates in synchrony with the cardiac cycle, in a state-dependent manner, providing, to our knowledge, the first evidence for cortical *in vivo* membrane potential state-dependent oscillation with the cardiac cycle.

One fascinating implication of our findings might be related to time perception. Indeed, recent evidence suggests that the subjective perception of time may rely on the processing of cardiac signals by the insular cortex.^35–38^ Here, we find that insular neurons fire at precise and specific timepoints within the cardiac cycle, a process that could support encoding of the internal, subjective passage of time by the insular cortex.^37,38^ Additionally, we show that pInsCtx neurons are preferentially tuned to the heartbeat at systole, which provides a strong mechanistical basis for the contraction of time perception when the heart contracts.^35^

Furthermore, we here find that this tuning is enhanced during emotion states. Interestingly, the systole has been associated with enhanced fear processing and detection in humans.^4,39^ In contrast, neural responses to external tactile stimuli in the motor cortex have been shown to be inhibited at systole.^40,41^ Taken together, these findings suggest a competition between interoceptive and exteroceptive signals during the different phases of the cardiac cycle.^42^

We further demonstrate that insular neurons are frequency-tuned to heartbeats. This is reminiscent of primary sensory cortices processing of exteroceptive information, such as the visual or auditory cortices. It would be interesting to explore the extent to which this precise interoceptive tuning resemble already described exteroceptive tuning. For example, frequency tuning in primary sensory cortices develops postnatally and is shaped during critical periods by the stochastics of the individual’s environment.^43,44^ While similar dependencies in the pInsCtx remain largely hypothetical at this point, they may carry broad impact and contribute to explain how bodily physiology shapes brain function and health.

Importantly, we report that cardiac signal processing is crucial for emotion state coding. Our study revealed that heartbeat tuning of pInsCtx neurons increased specifically during both aversive and positive emotion states. This finding might provide empirical evidence for classical and modern theories of emotions, such as the James-Lange theory or the constructed emotion theory.^45,46^

Additionally, we observed strong effects of metoprolol on cardiac reactivity to emotion and cardiac feature representation in the pInsCtx. While we cannot exclude that, additionally to its peripheral effects, metoprolol might act centrally as well, our findings highlight that heartbeat tuning is only affected during emotion. Moreover, metoprolol specifically disrupts insula reactivity and encoding of emotional cues, while it does not affect the encoding of sensory cues or overall firing rate of the insula. Overall, these results suggest that cardio-insular coupling supports emotion state coding. Moreover, these findings might help elucidate some of the mechanisms behind the known effects of beta-blockers against anxiety and acute psychological stressors.^47,48^

Taken together, our results show that cardiac signals are processed at a surprising accuracy and influence insula activity broadly. While the many potential functional implications of such cardioception in the insula and beyond are yet to be explored, we here demonstrate that they are crucial for one of the pInsCtx core functions, emotional state processing. Such processes might be altered in psychiatric disorders known to be associated with impaired interoception and insular activity such as anxiety disorders.^49,50^

## Material and Methods

### Experimental Animals

Animal experiments were conducted in accordance with the regulations from the government of Upper Bavaria. Male and female C57BL/6NRj mice received from the in-house breeding facility were used for this project. Animals were 1-6 months old, housed with littermates in groups of 3 and kept in an inversed 12h light/dark cycle. They were provided with ad libitum access to standard chow and water unless stated otherwise (see head-fixed appetitive conditioning).

### Surgical procedures

Thirty minutes before surgery, Metamizol (Hexal, 200 mg/kg bodyweight, s.c.) was applied for peri-operative analgesia. Anesthesia was initiated with 5% isoflurane in oxygen-enriched air and maintained at 1-2.5% throughout the surgical procedure. Mice were firmly secured in a stereotaxic frame (Model 51733D, Stoelting) and placed on a heating pad to maintain their body temperature at 37 °C. Eye ointment (Bepanthen, Bayer) was applied to protect the animal’s eyes. The head of the animals was shaved and lidocaine (Xylocain spray) was applied locally prior to incision.

### Headpost implantation

Custom-made stainless-steel head bars (35x3x3 mm) were affixed to the skulls of mice using dental cement (Super Bond, Génerique International) while under isoflurane anesthesia. The skull underneath the head implant opening was covered with dental cement and future locations for the craniotomies above pInsCtx (from Bregma: AP: -0.40 mm, ML: ± 4.10 mm) or wM1 (from Bregma: AP: 1.0 mm, ML: ± 1.0 mm) were marked.

### Heart rate electrode implantation

Surgical staples (FST 12040-01, Fine Science Tools) were secured onto the shaved skin on both lateral sides of the abdomen and on the back of the neck using suture clip forceps (FST 12018-12, Fine Science Tools).

### Craniotomy

The day of recording, animals were anesthetized using isoflurane and a craniotomy (approx. 1.5 mm diameter) was drilled at the previously marked location. The dura was carefully removed above the future recording site and the craniotomy was covered with silicon elastomer (Kwik-cast, World Precision Instruments). Mice were placed in a recovery cage for at least 1.5 hours before recording. Mice were typically recorded over two consecutive days.

### Electrophysiological recordings

#### In vivo whole-cell patch clamp recordings

Recordings were performed as previously described.^51,52^ Briefly, electrodes (BF150-86-10, Science Products) were prepared with a vertical puller (PC-10, Narishige) to achieve 4-6 MΩ tip resistance. The recording depth for the insular cortex was obtained from stereotaxic coordinates, and recording location was marked at the end of the recording either with biocytin (Sigma-Aldrich, B4261) or DiI (Life Technologies GmbH, V22885). Recording electrodes were filled with a solution containing (in mM) K-gluconate 135; KCl 4; phosphocreatine 10; MgATP 4; Na3GTP 0.3; HEPES 10.^53^ Whole-cell recordings were obtained using the blind-patch approach.^51,54^ Access resistance and membrane capacitance of the cells were monitored on-line and analyzed offline. The mean access resistance was 51.8 ± 2.4 MΩ. The membrane potential was not compensated for the liquid junction potential. Recordings were not analyzed if the assess resistance exceeded 100 MΩ or when the action potential peak dropped below 0 mV.

#### Silicon probe recordings

The silicon probe was first coated in 10% DiI diluted in 96% ethanol for later visualization of the probe track. The probes were connected to the headstage and securely mounted to a 3-axis micromanipulator (S-IVM-3000C, Scientifica). Mice were head-fixed to the recording setup, and the silicon elastomer was removed using forceps to expose the craniotomy. The silicon probe was lowered until touching the brain surface and the depth coordinates were set to 0. The probe was then inserted at a speed of approximately 1µm/sec, as visualized with the LinLab software (Scientifica). The recording depth was set at -3.5mm, determined empirically in tests experiments prior to the recordings. Once the target depth was reached, a minimum waiting period of 15 minutes was observed before starting the recordings to allow the tissue to recover from the penetration.

### Behavioral conditioning

#### Training and habituation to head-fixation

After headbar implantation, mice were monitored and weighted every day for 5 days. Following recovery, mice were progressively habituated to head-fixation: 10 min on day 1, 30 min on day 2, 60 min on day 3, and up to 120 min (for whole-cell recordings) on day 4. Thereafter, mice were head-fixed 60 or 120 min each day until they fully habituated and showed a comfortable posture (no contortion of the body, calm behavior alternating between running and immobility). Typically, mice habituated within 2 to 10 days. During the final days of training, mice were head-fixed on the wheel used for the recordings and habituated to the ambient light and noises that would occur during an experiment.

#### Head-fixed appetitive conditioning

After recovery and wheel habituation, mice underwent appetitive conditioning on the wheel. A day prior, water restriction started (water bottles taken out of the cage), and 2 ml of water was delivered every day to a dish placed within each cage. Mice were weighted and handled daily. This behavioral pavlovian conditioning was adapted from a published protocol.^55^ On the first day, mice were head-fixed and exposed to unpredictable drops of sucrose (10% sucrose in water; 2.0–2.5 μl) delivered intermittently for one hour (approximately 50 drops per hour) through a gravity-driven, solenoid-controlled lick tube. This step allows the animals to learn to lick the spout, and drink the sucrose drop. This behavior was acquired typically on day1. After this sucrose habituation step, mice underwent pavlovian appetitive conditioning. During each conditioning session, two cues (CS+: 3 kHz 50ms tones CS-:2 kHz constant tones, 2 s) were randomly presented 50 times before the delivery of sucrose (CS+, 10% sucrose in water; 2.0–2.5 μ l) or no sucrose (CS−), such that there was a one second trace interval between delivery of the CS+ and sucrose. The inter-trial interval between the previous reward delivery (CS+) or withholding time (CS−) and the next cue was chosen as a random sample from a uniform distribution bounded by 40s and 80s.

After 6-16 days of conditioning, two consecutive days of pInsCtx recordings were performed. During these recordings, mice were exposed to 10 randomly presented CS- and CS+, as well as a new CS (neutral CS+ on day1/fear CS+ on day2; 500ms x 4 upsweeps ranging from 7kHz to 20kHz).

#### Aversive conditioning

In experiments where mice underwent appetitive conditioning, they were also subjected to an auditory fear conditioning paradigm. This allowed us to compare appetitive and aversive neuronal, behavioral and physiological responses in the same animals. Briefly, on day 1 after the first electrophysiological recording session, mice were placed in a commercial fear conditioning setup (Ugo Basile, Italy), consisting of a behavior box, an electric grid floor, and a light source located in a soundproof chamber. The chambers were additionally equipped with electrostatic speakers for high precision sound delivery (Tucker Davis Technologies). The last second of the neutral CS+ was paired 5 times with a 0.4mA footshock (unconditioned stimulus, US), in order to create an aversive memory associated with this CS+ presentation. As a readout of an animal’s fear response, freezing behavior (defined as complete immobility with the exception of breathing) was analyzed. Freezing was automatically quantified using the software ANYmaze (Stoelting). Behavior of the mice was videotaped from the side of the chamber and the software calculated a “freezing score” depending on the number of pixel changes between frames. If the freezing score fell below an empirically determined threshold for at least 2 s, mice were considered to be freezing.

### In vivo pharmacology

#### Pharmacological manipulation of heart rate

In a subset of experiments, we manipulated heart rate to investigate the impact of a reduced heart rate on pInsCtx activity and encoding of emotions. To do so, on day 2 of the electrophysiological recordings, mice received an intra-peritoneal injection of the beta-blocker metoprolol (3mg/kg) after a baseline recording. Metoprolol is a selective antagonist of the β1-adrenergic receptor found primarily in cardiac tissues and has no intrinsic sympathomimetic activity. By competing for the biding sites with adrenaline and noradrenaline, metoprolol blocks their activation which leads to a reduction in heart rate. Metoprolol is known to only be moderately lipophilic, which leads to a lower blood-brain barrier crossing than other commercial β-blockers such as propranolol.^56^

#### Inhibition of synaptic transmission in the pInsCtx

In a subset of experiments, we aimed to investigate if synaptic transmission was necessary for heartbeat tuning of pInsCtx neurons. To answer the question, we inhibited synaptic transmission locally in the pInsCtx by infusing a cocktail of synaptic inhibitors: APV 2mM, CNQX 1mM and mecamylamine 1mM. APV is a selective antagonist of NMDA receptors and inhibits their glutamate biding site. CNQX is an AMPA/kainate glutamate receptors antagonist. Mecamylamine is an antagonist of nicotinic acetylcholine receptors. The synaptic blockers were diluted in Phosphate Buffer Saline (PBS 1X prepared from a 10X stock solution) and Alexa Fluor 594 was added to the solution for post-hoc visualization of the drug infusion site. On the day of the electrophysiological recordings, the solution was loaded into a glass electrode connected to a picospritzer which allows for pressure ejection of small volumes. The glass electrode was inserted at a 20 degrees angle from the brain surface, at the following coordinates from Bregma: +0.5 AP, 4 ML, -2.5 DV. These coordinates were obtained using Pinpoint (https://github.com/VirtualBrainLab/Pinpoint). Recordings were conducted as follow: baseline activity recordings (> 15min) followed by a drug infusion consisting of 2 pulses of 500ms at 1Hz repeated every 4 minutes. In some recordings, only one drug infusion was performed in order to observe activity returning back to baseline and estimate to time for washing off the drugs.

### Infrared videography for facial and locomotion monitoring

We acquired data at 30 Hz using a USB 3.0 monochrome camera (BFS-U3-13Y3M-C, Point Grey Research), positioned perpendicularly to the left side of the mouse’s head, capturing both the face and the front paws on the wheel for locomotion detection. Illumination was provided by a battery-operated IR light (Andoer Mini IR Night Vision Light).

### Histology and microscopy

At the end of the electrophysiological recordings, animals were anesthetized with a mixture of Ketamine and Xylazine (100 mg/kg and 20 mg/kg bodyweight, respectively, Serumwerk Bernburg) and perfused intra-cardially with 4% paraformaldehyde (PFA) in phosphate buffered saline (PBS). Brains were post-fixed for an additional 24 h in 4% PFA at 4 °C. Then, coronal sections were cut at 70 μm with a VT1000S vibratome (Leica Biosystems). For biocytin visualization in the patch-clamp experiments, slices were incubated overnight at 4 °C in 1:1000 Alexa Fluor 488 conjugated in the same solution. Sections were mounted onto the glass slides in anatomical order using Vectashield Antifade Mounting Medium with DAPI (Vector Laboratories). Images of the brain sections were acquired using a fluorescence microscope (VS120, Olympus).

### Data acquisition, processing and statistical analysis

#### In vivo whole-cell patch clamp recordings

All recordings were made in the current-clamp configuration. Recordings were obtained with a Multiclamp 700B amplifier connected to the Digidata 1440A system digitizing analog input channels at a 16 bits resolution and at up to 250 kHz each. Data were acquired with pClamp 10 (Molecular Devices), digitized at 20 kHz, filtered at 3 kHz, and analyzed offline with Clampfit 10.4 (Molecular Devices) and custom pipelines in Python (Anaconda, distribution for Python version 3.12) which made use of the SciPy^57^ and Neo^58^ packages.

#### In vivo silicon probe recordings

Recordings were performed using Acute 64 channels 2 shanks (H2 and H6) silicon probes from Cambridge Neurotech, using the Digital Lynx SX (NeuraLynx) interface board, sampled at 20 kHz. Amplification and digitization were done on the head stage. The recordings were performed using the Cheetah software (NeuraLynx). Initial clustering was performed using Kilosort 2.5,^59^ before manual curation using Phy (https://github.com/cortex-lab/phy/).

#### Pupil size and locomotion

The area of the pupil and the motion energy change around the paw area during locomotion was determined offline using FaceMap.^60^ An ROI was drawn manually around the pupil of the animal. The pupil area was defined as the area of a Gaussian fit on thresholded pupil frames, where pixels outside the pupil were set to zero.

#### Electrocardiogram recordings (ECG) and heart rate processing

ECG was recorded using one of the two patch clamp headstages. To be able to record voltages changes associated with cardiac activity, recordings were made in the current-clamp configuration. Briefly, the two surgical staples on both sides of the abdomen were clamped using alligator clips with wires soldered together to a gold female pin connected to the recording input of the headstage. The ground electrode placed on the neck was clamped independently and connected to a gold pin to the ground input of the headstage. Recordings were obtained with a Multiclamp 700B amplifier connected to the Digidata 1440A system digitizing analog input channels at a 16 bits resolution and at up to 250 kHz each. Data was acquired with pClamp 10 (Molecular Devices) and digitized at 20 kHz. The data was then processed using custom Python pipelines. The ECG trace was first down sampled to 10kHz and smoothed using a Savitzky–Golay filter. R peaks were detected by extracting when the derivative dV/dt signal crossed a threshold that was adjusted depending on the signal/noise ratio. Each ECG trace was inspected manually and the R peak detection was adjusted if needed to make sure no heartbeat was either missing or falsely detected. Instantaneous heart rate in beats/min was then computed by counting R peaks in 1-second sliding windows.

#### Heart rate variability

Heart rate variability was calculated as the root mean squared of successive differences between consecutive heartbeats (RMSSD), in ms. The RMSSD is obtained by first calculating each successive time difference between heartbeats in ms. Then, each of the value is squared and the result is averaged using a rolling mean window of 30 consecutive datapoints. Finally, the square root of the average values is calculated. The RMSSD reflects the beat-to-beat variance in heart rate and is the primary time-domain measure used to estimate vagal tone.^61^

#### Heartbeat tuning

To determine if a neuron was heartbeat tuned, we first calculated the phase of the units’ firing around heartbeat. To do so, we detected the two nearest heartbeats for each spike time, and computed where the spike occurred within that heartbeat interval, normalized to a 0 to 2π phase in radian. When all the phases were obtained for a unit, we determined if the phases were non-uniformly distributed, i.e. if spikes are synchronized to the heartbeats, using the Rayleigh test, a circular statistics method.^62^ If the Rayleigh test returned a p-value below 0.05, we classified the unit as heartbeat tuned, as this suggests significant phase locking between a neurons’ firing and heartbeats.^11^

To determine a neurons’ preference for tuning to heartbeat at specific frequencies, we analyzed heartbeat tuning over time. We first split the data into time bins (from 200ms to 10s), calculated the phases of spiking relative to the nearby heartbeats, and then applied the Rayleigh test to compute if spikes are phase-locked to heartbeat within that time bin. This allowed us to identify when a neuron was heartbeat tuned across time. We then extracted the heartrate values in Hz during significant heartbeat tuning for each neuron. Finally, we determined tuning preferences for each neuron by detecting the peaks in the heartrates at which a neuron would tune in. We classified neurons has having a single, two, or more frequencies tuning preference if the heartrate at which they would tune in showed single peak, two peaks, or more peaks, respectively.

#### Correlations between pInsCtx dynamics and heart rate, pupil and locomotion

In order to address how strongly pInsCtx activity co-varied with heart rate, pupil, or movement, we calculated the correlations between pInsCtx firing rates or pInsCtx membrane potential and heart rate, pupil size, and locomotion. We first z-scored each time series before computing the Pearson correlation between each pair of variables, either neuron-by-neuron using firing rate or membrane potential, or average population activity using the average pInsCtx across a recording session.

#### Random spike trains generation

To generate random spike trains, we randomized the real spike trains of pInsCtx using the “spike train surrogates” function of the elephant package (https://zenodo.org/records/13133971). Specifically, the spikes were randomly repositioned inside the time interval, keeping the spike count, and generating a Poisson spike train with time-stationary firing rate. We generated 14 recording sessions with the same number of units than our real pInsCtx recordings, for comparison.

#### Variable decoding from pInsCtx activity

In order to assess how pInsCtx encodes specific behaviors and physiological states, we performed decoding of heart rate, pupil size, and locomotion from pInsCtx neural activity using Lasso regression. Lasso is a regularized L1-linear regression model predicting an outcome variable from input features while selecting the most important predictors.^63^ We used a 5 splits time series cross-validator to provide train and test data. We first z-scored each time series for standardization. Decoding accuracy was computed as 1 – MSE (Mean Squared Error) between the observed and the predicted variable.

#### Detection of heart rate increase

We identified heart rate increases and categorized them based on whether they occur during or outside locomotion events. We first z-scored heart rate and detected increases as the timepoints where the z-scored heart rate went above 0. We only kept events lasting more than 2 seconds. We then computed the firing rates of each pInsCtx unit relative to all increases in heart rate during locomotion or immobility. For each unit, we represented the average firing rate relative to heart rate increases as a heatmap. We classified the units as non-responsive, activated or inhibited during heart rate increases by comparing their firing rates 2s before versus 2s after the heart rate increase, using t-test on related samples. A neuron was classify as significantly modulated if the p-value was lower than 0.05. Neurons were further classified as inhibited or activated depending on the sign of the modulation.

#### Freezing detection

We detected freezing episodes using a similar method as we previously described.^18^ We detected pupil dilations occurring during periods of complete immobility and selected the dilations that lasted minimum 2s.

#### PCA

We used Principal Component Analysis (PCA) as a dimensionality reduction technique to visualize neural activity patterns across different events. We first extracted all neuronal activity across trials for every animal, and z-scored them. We performed PCA (15 components) and visualized the first two principal components PC1 and PC2 previously smoothed using a Gaussian filter (sigma=3). We visualized the components using confidence ellipses showing the spread of neural activity for each trial type in the PC space.

#### Linear Classifier

To address if CS identities are encoded by pInsCtx population activity in control animals versus metoprolol treated animals, we used a linear-kernel support vector machine (SVM) classifier.^64^ Firing rates were computed in 50ms bins for a -2s to +10s window relative to CS onsets, for each mouse. For each time bin, we trained an SVM to classify the trials based on pInsCtx neural activity, using a 5-fold cross-validation. We then computed accuracy scores across folds, as the fraction of the correct predictions for a given fold. After the 5 folds were evaluated, we averaged the 5 scores for each time bin to get the overall decoding performance at that time.

We then determined whether the decoding accuracy was significantly above chance level (0.5) for distinguishing between events, both before and after metoprolol. To do so, we performed permutation tests by comparing the real accuracies to a null distribution generated by shuffling labels. Briefly, for each time bin, labels were shuffled randomly and the SVM accuracy computed for the shuffled labels. This step was repeated 1000 times to build a distribution of chance-level accuracies. Then, we calculated the fraction of null accuracies > real accuracies. Low p-values indicates that the real accuracies are unlikely to be due to chance. Finally, we applied a false discovery rate (FDR) correction to account for multiple comparisons, using the Benjamini-Hochberg procedure.

### Statistics

Statistical analysis was performed using Python. Two-group single variable comparisons were performed using non-parametric Wilcoxon tests or parametric Student t-tests, depending on the normality of the data. Correlation analyses were performed by calculating Pearson’s correlation coefficient. Group comparisons were made using Friedman tests (non-parametric) two-way ANOVA (parametric) followed by post hoc tests (Wilcoxon or Bonferroni) comparing the groups in case the main effect was statistically significant (p < 0.05). Detailed information about the type and results of all statistical procedures can be found in figure legends. All animal numbers are reported in Figures and their legends.

## Data availability

All data and code will be made available on public repositories upon publication.

## Acknowledgements

We would like to thank Cornelia Flachskamm, Rainer Stoffel for help with wheel training of the animals. Mario Carta, Dennis Nestvogel, Oriane Mauger and Anna Zych for feedback on earlier versions of the manuscript, and the Gogolla group for feedback and discussions throughout the years. Dennis Nestvogel for providing suggestions throughout the project. Eunjae Cho for technical help, Alexandra Klein and Nate Dolensek for technical assistance during the initial phases of the project. This project was supported by the Max Planck Society with generous core funding to N.G., the European Research Council (ERC) under the European Union’s Horizon 2020 research and innovation program (ERC-2017-STG, grant agreement n° 758448 to N.G.).

## Author contributions

M.M. and N.G. conceptualized and supervised the project. M.M. designed, performed and analyzed all experiments. M.M. and N.G. interpreted the results. J.Y. helped with initial implementation of silicon probes recordings and analysis. A.R. and B.S. performed histological controls.

M.M. wrote the original draft. M.M. and N.G. edited and contributed to the final version of the manuscript.

N.G. acquired funding.

