## Supplementary Figures 1-10 for "Cardiac signals shape insular cortex activity and emotion coding"

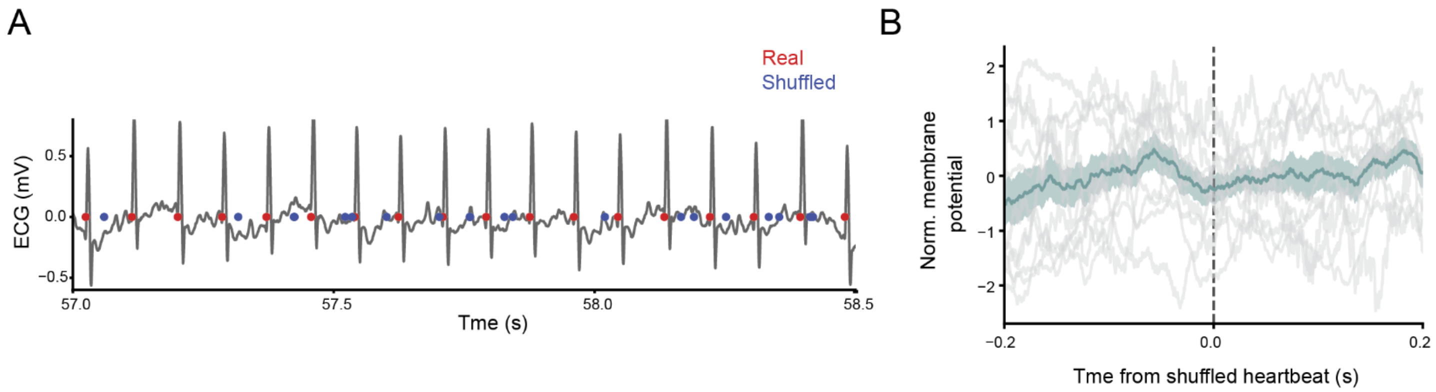


**Extended Data Fig.1 (related to main Fig.1). pInsCtx membrane potential fluctuations around shuffled heartbeats**

**(A)** Example trace showing real heartbeats detection (red) and shuffled heartbeats (blue).

**(B)** Membrane potential fluctuations averaged relative to shuffled heartbeats. Data is shown for individual mice (grey) and average ± s.e.m. of 10 cells in 7 mice (green).

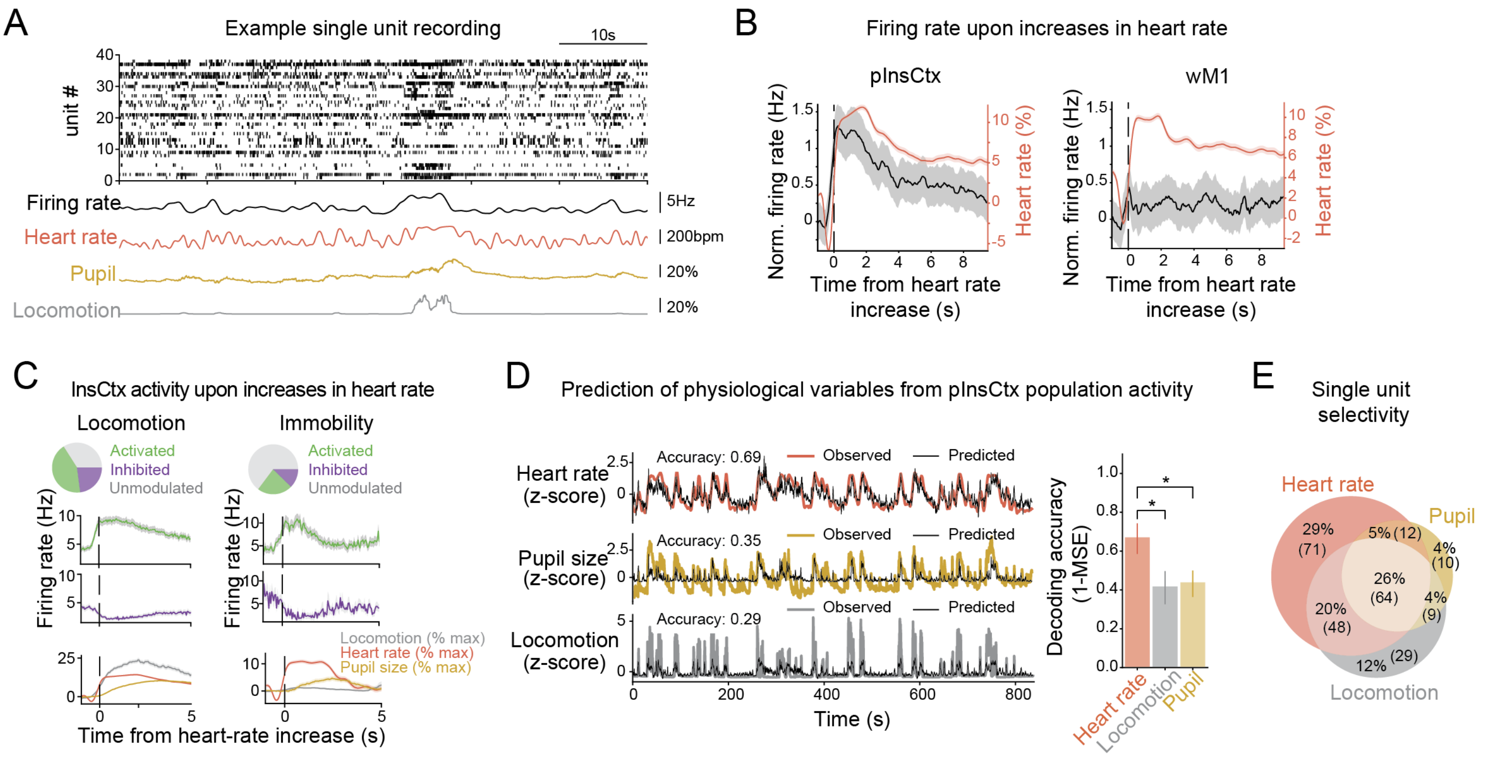


**Extended Data Fig.2 (related to main Fig2). Activity in the pInsCtx is predominantly influenced by cardiac signals**

**(A)** Example silicon probe pInsCtx recording: raster plot where each line represents one pInsCtx unit and each tick corresponds to an action potential (top), the average corresponding firing rate (black), heart rate (red), pupil size (yellow) and locomotion (grey).

**(B)** Firing responses of neurons in the pInsCtx (left, black) versus the primary whisker motor (wM1) cortex (right, black) relative to increases in heart rate (red). pInsCtx: n=10 mice, wM1: n=6 mice. Traces represent mean ± s.e.m.

**(C)** pInsCtx activity upon heart rate increases during locomotion (left) versus immobility epochs (right). Neurons were separated into significantly activated by heart rate increases (green), inhibited (purple), or unmodulated (grey) locomotion (student t-test, *p<0.05). During locomotion (left), 43.1% of all neurons were activated, 22.9% were inhibited and 34.0% were not modulated when heart rate increased (top left pie chart). During immobility (right), 22.9% neurons were activated, 12.3% inhibited and 64,8% were not modulated when heart rate increased (top right pie chart). Traces represent mean ± s.e.m.

**(D)** Left: Example traces of a linear, L1-regularized model trained on pInsCtx population activity to predict heart rate (top), pupil dynamics (middle) or locomotion (bottom). Z-scored observed heart rate (red, top), pupil (yellow, middle) and locomotion (grey, bottom) overlaid with their corresponding prediction in black. Right: Quantification of the decoding accuracy of the linear models expressed as 1 - Mean Squared Error (decoded behavior - true behavior) (Friedman test (p<0.05) followed by post hoc Wilcoxon test, *p < 0.05). Bars represents mean ± s.e.m. for n=10 mice.

**(E)** Proportion of pInsCtx neurons selective for heart rate increases (red), pupil increases (yellow) or locomotion increases (grey).

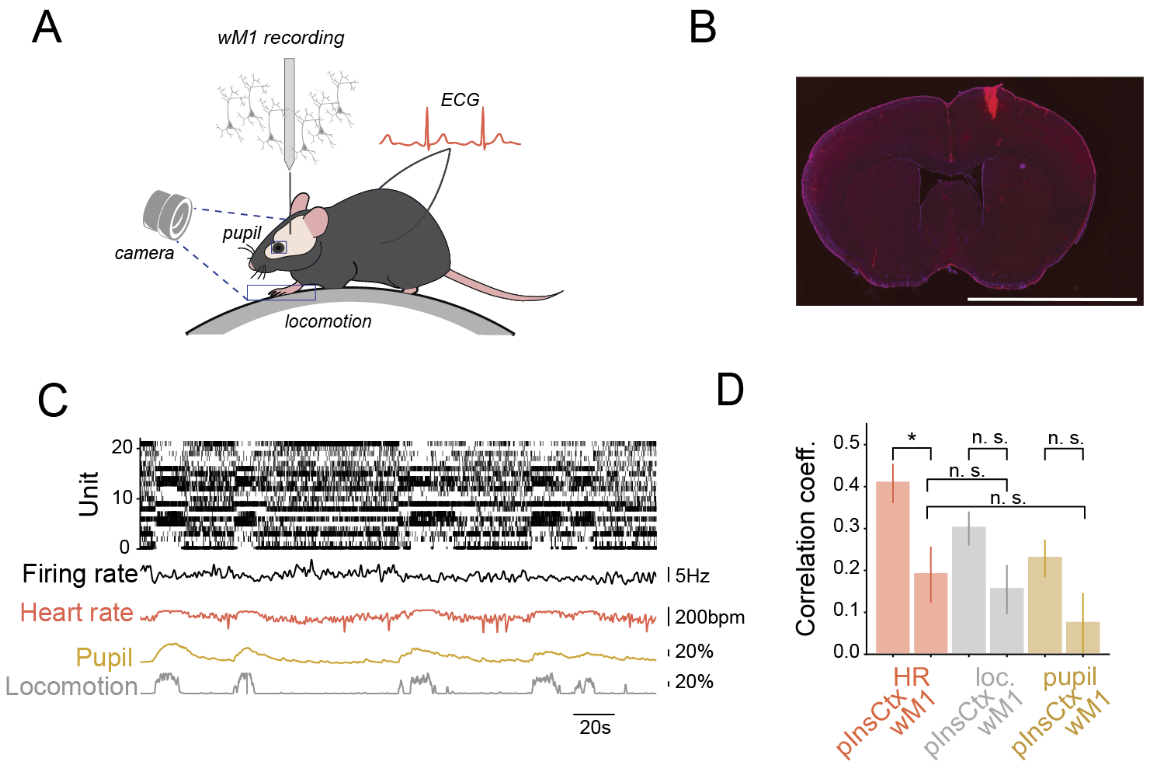


**Extended Data Fig.3 (related to main Fig2). Motor cortex activity is only weakly modulated by cardiac signals**

**(A)** Schematic illustrating silicon probe recordings in the wM1 of head-fixed mice.

**(B)** Slide scanner image of a coronal section a mouse brain, showing the recording electrode track in wM1. Bregma level +0.6. Scalebar 5mm.

**(C)** Example wM1 recording: (top) raster plot with each black dot corresponding to an action potential, average corresponding firing rate (black), heart rate (red), pupil size (yellow) and locomotion (grey).

**(D)** Pearson's correlation coefficients of pInsCtx population average firing rate to heart rate, pupil or locomotion in pInsCtx versus wM1 (Wilcoxon test, $*p < 0.05$). (n=14 recordings in N=10 Mice for pInsCtx; n=10 recordings in N=6 mice for wM1). Bars represents median ± s.e.m.

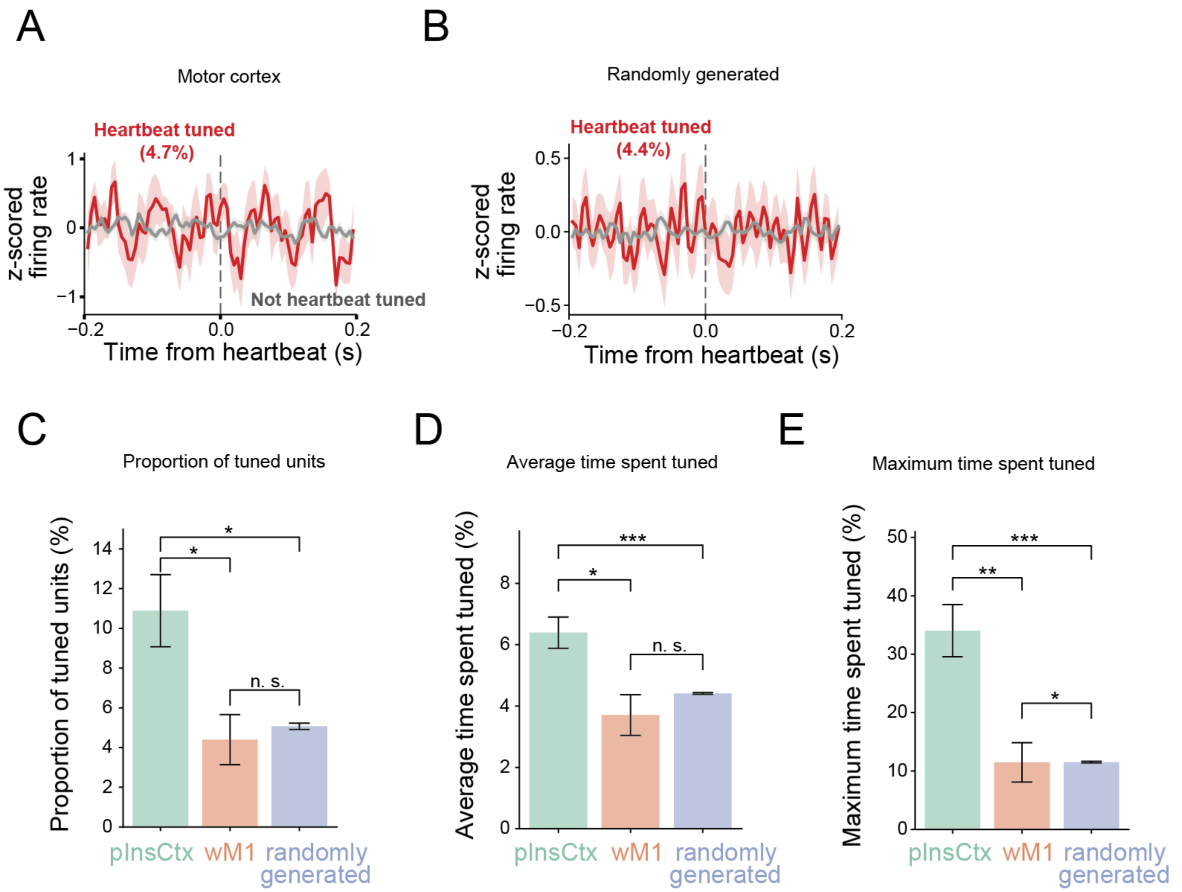


**Extended Data Fig.4 (related to main Fig.2). Heartbeat tuning in motor cortex and randomly generated spike trains**

**(A)** Firing rate relative to heartbeats for all significantly heartbeat tuned neurons in the motor cortex (rayleigh test, *p<0.05; 4.7%, n=8 units in N=10 recording sessions in 6 mice) compared to untuned neurons.

**(B)** Firing rate relative to heartbeats for all significantly heartbeat tuned neurons in randomly generated recordings (rayleigh test, *p<0.05; 4.4%, n=23/524 units in N=14 randomly generated recordings) compared to untuned neurons.

**(C)** Proportion of heartbeat tuned units in pInsCtx (green), motor cortex (orange) and randomly generated recordings (blue). (*p<0.05, Mann–Whitney U test).

**(D)** Average time spent heartbeat tuned in pInsCtx (green), motor cortex (orange) and randomly generated recordings (blue) (*<p0.05, ***p<0.001, Mann–Whitney U test).

**(E)** Maximum time spent heartbeat tuned in pInsCtx (green), motor cortex (orange) and randomly generated recordings (blue) (*p<0.05, **p<0.005, ***p<0.001, Mann–Whitney U test).

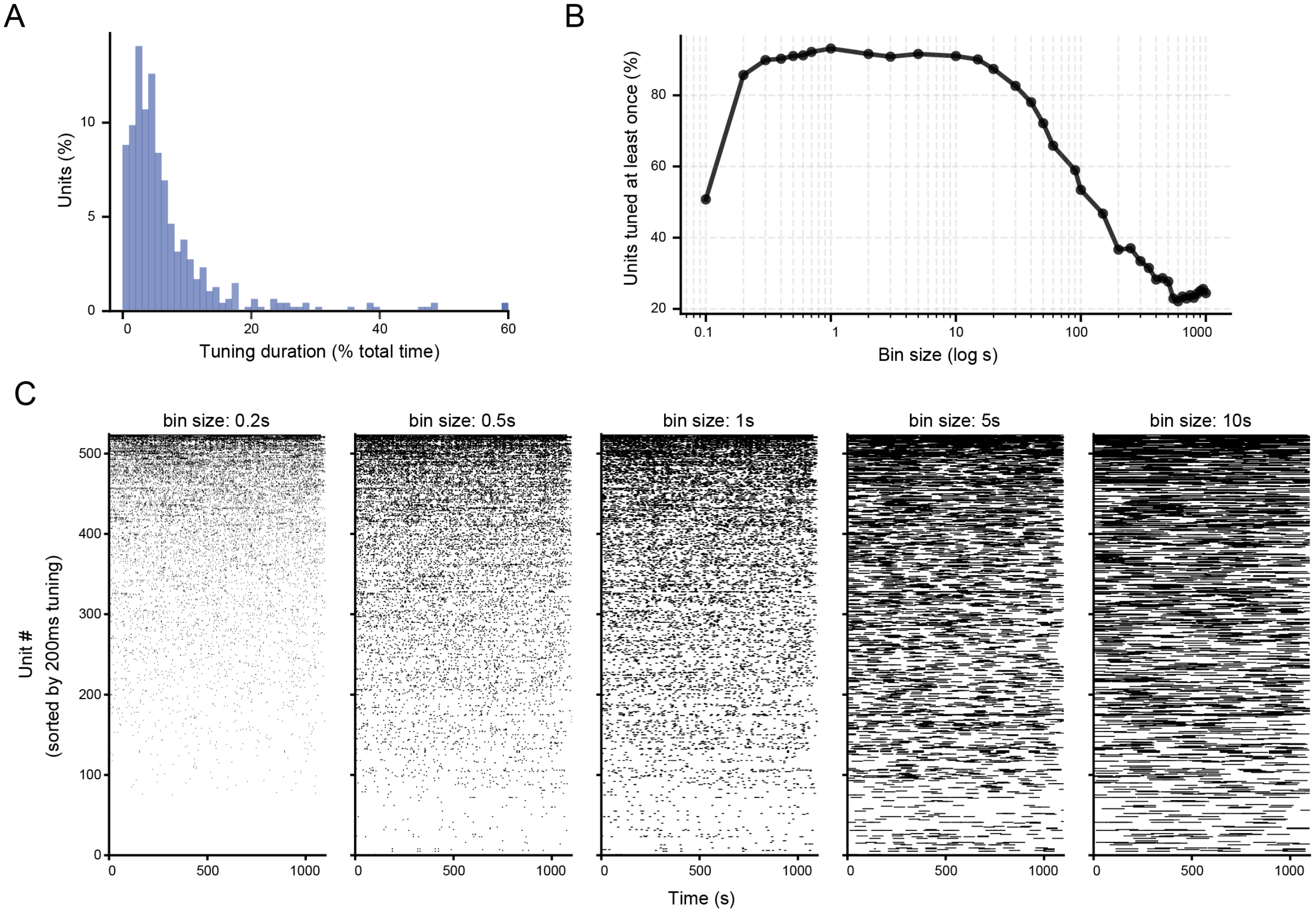


**Extended Data Fig.5 (related to main Fig2). Heartbeat tuning characterization**

**(A)** Percentage of unit heartbeat tuned at least once depending on tuning timebin.

**(B)** Distribution of time spent tuned as a percentage of the whole recording session, for timebins of 10s.

**(C)** Scatter plots showing heartbeat tuning events of all recorded units for different timebins, sorted according to the total time spent tuned for 200ms timebin. Note that the general sorting is preserved, with highly tuned units clustering towards the top and low tuned units clustering towards the bottom, suggesting that timebin size has a low impact on the heartbeat tuning analysis.

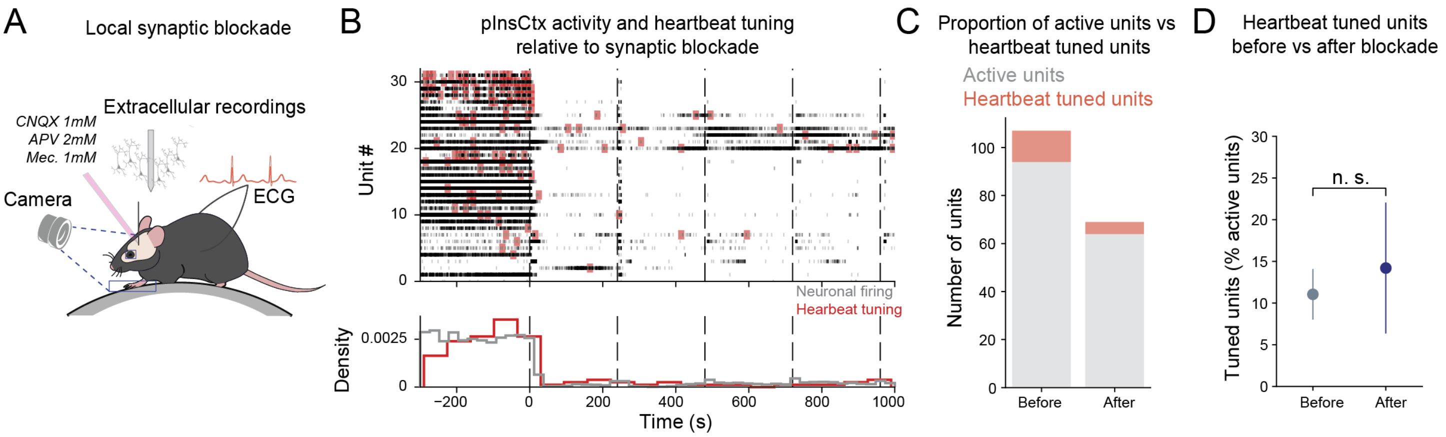


**Extended Data Fig.6 (related to main Fig2). Heartbeat tuning requires synaptic transmission**

**(A)** Schematic of silicon probe recordings in the pInsCtx of head-fixed mice with synaptic blockers infusion using a glass electrode filled with CNQX 1mM, APV 2mM and Mecamylamine 1mM in PBS with Alexa.

**(B)** Recording with synaptic blockers infusion. (top) Raster plot of recorded units activity (black) and heartbeat tuning (red) relative to synaptic blockers infusion (dashed lines). (bottom) density histogram of firing rates and heartbeat tuning relative to synaptic blockers infusion (dashed lines).

**(C)** Proportion of heartbeat tuned neurons among the active population (firing rate > 0.5Hz) before and after synaptic blockers infusion. 13/107 heartbeat tuned units before vs 5/69 heartbeat tuned units after synaptic blockers infusion. N=5 mice.

**(D)** Percentage of heartbeat tuned pInsCtx units during before (grey) vs after (blue) synaptic blockade infusion (n.s., student t-test). Data represents mean variable ± s.e.m. N=5 mice.

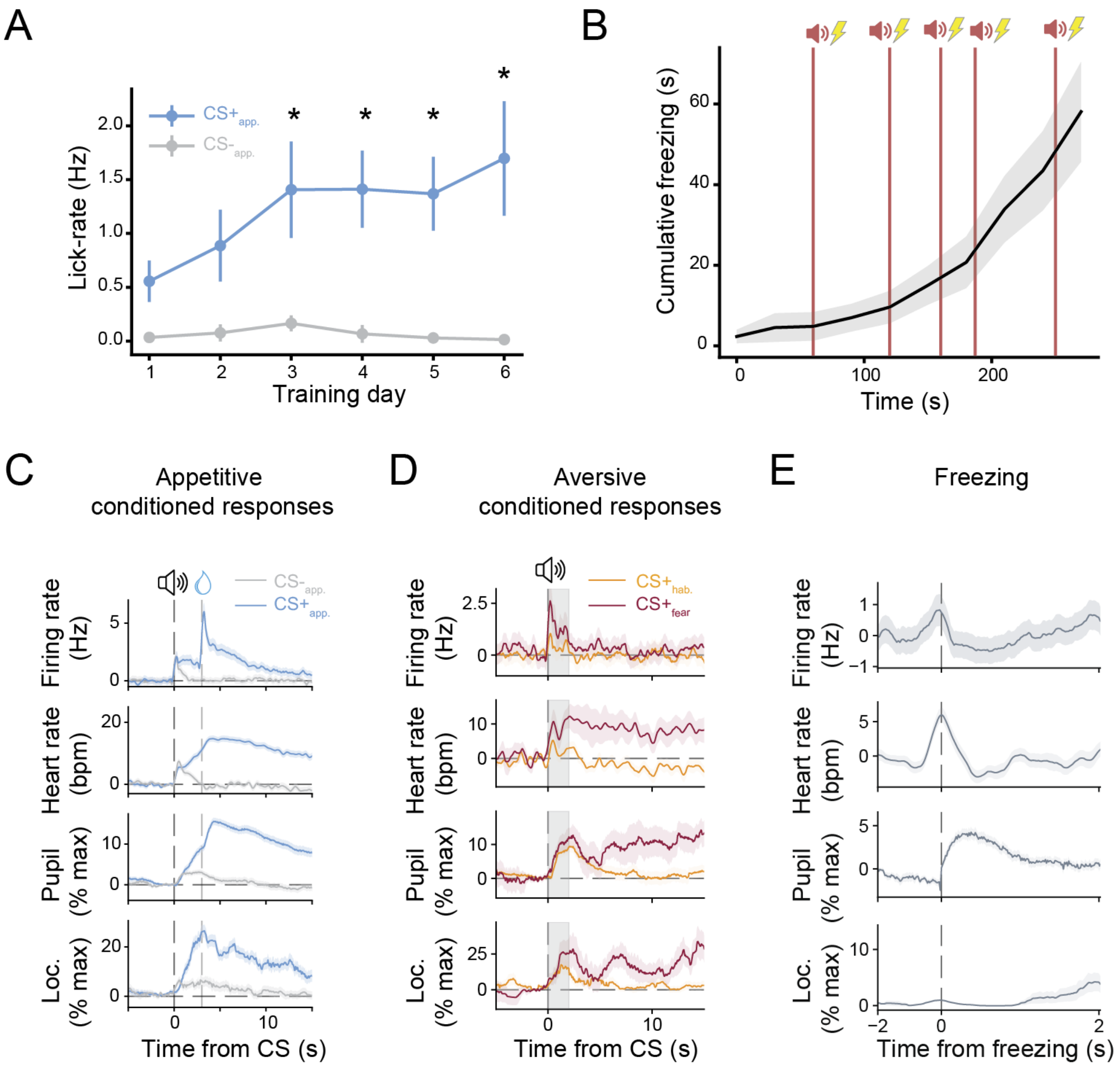


**Extended Data Fig.7 (related to main Fig.3). Changes in pInsCtx activity and behavior during emotion states**

**(A)** Lick-rate in Hz for CS- and CS+ across training days. The lick-rate change is computed as the difference between the number of licks/second 1s before versus 3s after CS onset. (*p<0.05, paired student t-test FDR corrected). N=6 mice.

**(B)** Cumulative freezing in seconds during fear conditioning. The footshock occurs during the last second of the CS+. N=10 mice.

**(C)** Conditioned responses to $CS+_{app.}$. Normalized pInsCtx firing rate, heart rate, pupil size, locomotion, relative to the start of $CS-_{app.}$ (grey) and $CS+_{app.}$ (blue). Traces are normalized by the activity 2s before the CS presentation. Sucrose is delivered 3s after the CS+ starts and is indicated by the second dashed line. Data represents mean variable ± s.e.m. N=10 mice per group.

**(D)** Conditioned responses to aversive CS. Normalized pInsCtx firing rate, heart rate, pupil size, and locomotion relative to the start of $CS+_{hab}$ (day1, orange) or the $CS+_{fear}$ (day2, red). Data represents mean variable ± s.e.m. N=4 mice per group.

**(E)** Freezing related changes in pInsCtx firing rate, heart rate, pupil size, and locomotion relative to the start of freezing on day2 after fear conditioning. Data represents mean variable ± s.e.m. N=4 mice.

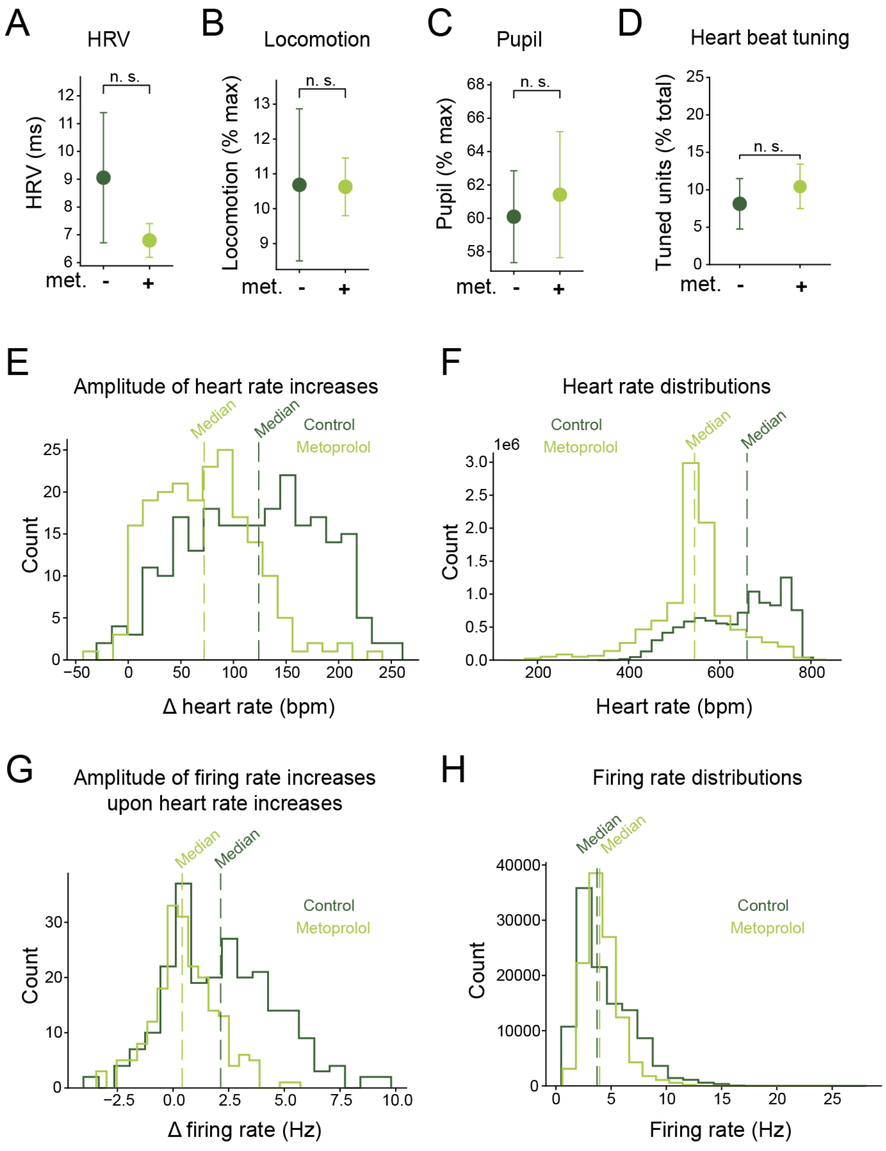


**Extended Data Fig.8 (related to main Fig.4). Metoprolol effects on physiological variables and pInsCtx activity**

**(A)** Heart rate variability (RMSSD, ms) before and after metoprolol administration. Data represents mean ± s.e.m. N=4 mice.

**(B)** Average locomotion before and after metoprolol administration. Data represents mean ± s.e.m. N=4 mice.

**(C)** Average pupil size before and after metoprolol administration. Data represents mean ± s.e.m. N=4 mice.

**(D)** Percentage of heartbeat tuned units before and after metoprolol administration. Data represents mean ± s.e.m. N=4 mice.

**(E)** Distribution of the amplitude of increases in heart rate in control (dark green) and metoprolol (light green) treated mice. N=4 mice per group. Dashed lines represent median increase in heart rate.

**(F)** Distribution of all heart rate values in control (dark green) and metoprolol (light green) treated mice. N=4 mice per group. Dashed lines represent median heart rate.

**(G)** Distribution of the amplitude of increases in pInsCtx firing rate upon heart rate increases in control (dark green) and metoprolol (light green) treated mice. N=4 mice per group. Dashed lines represent median increase in firing rate.

**(H)** Distribution of all pInsCtx firing rate values in control (dark green) and metoprolol (light green) treated mice. N=4 mice per group. Dashed lines represent median firing rate.

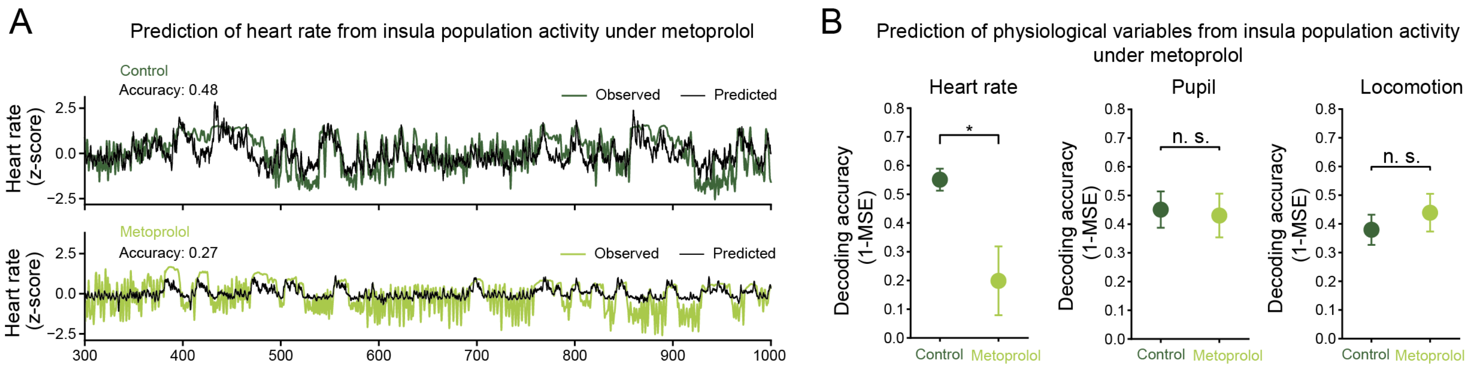


**Extended Data Fig.9 (related to main Fig.4). Metoprolol impairs heart rate prediction from pInsCtx activity**

**(A)** Top: Example from one control mouse on day 1 where a linear, L1 regularized model was trained on pInsCtx population activity to predict heart rate. Observed heart rate in dark green overlaid with the prediction in black.

Bottom: Same as (top) for the same mouse treated with metoprolol on day 2. Observed heart rate in light green overlaid with the prediction in black.

**(B)** Quantification of the decoding accuracy of the linear models expressed as 1 – Mean Squared Error (decoded behavior – true behavior) for (left) heart rate, (middle) pupil, and (right) locomotion (Mann-Whitney-U test, *$p < 0.05$). Data is presented as mean ± s.e.m. N = 4 mice per group.

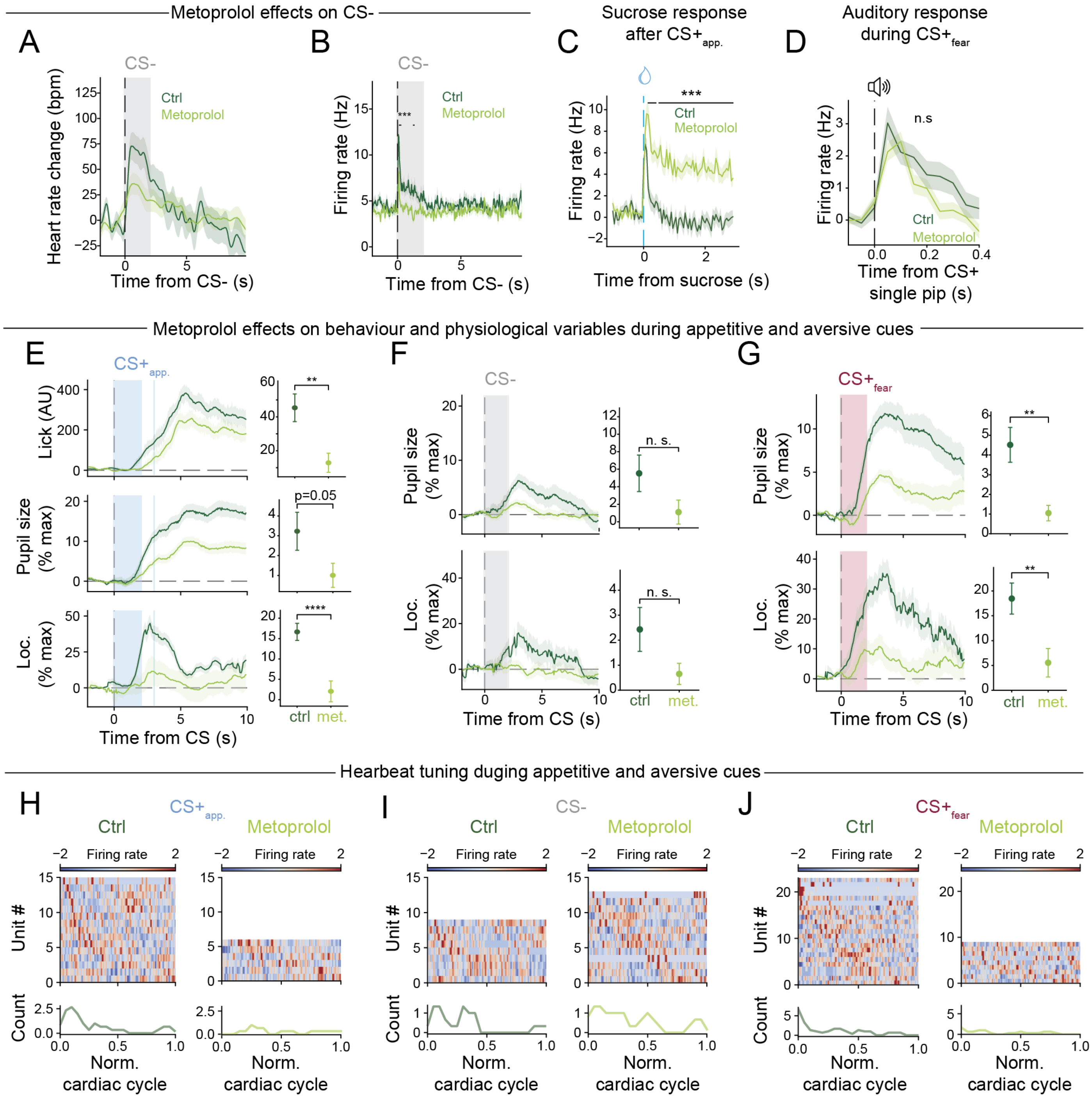


**Extended Data Fig.10 (related to main Fig.4). Metoprolol effects on physiological variables and pInsCtx activity during emotions**

**(A)** Heart rate change relative to the CS-$_{app.}$ in control (day1) versus metoprolol-treated (day2) animals. Traces are normalized to 2s before the start of the CS. (n.s., repeated measures ANOVA). N=4 mice per group. Data represents mean heart rate ± s.e.m.

**(B)** Firing rate of pInsCtx neurons relative to CS-$_{app.}$ in control versus metoprolol-treated animals (***p<0.001, repeated measures ANOVA with Bonferroni correction). N=4 mice per group. Data represents mean firing rate ± s.e.m.

**(C)** Firing rate of pInsCtx neurons relative to the sucrose delivery in control versus metoprolol-treated animals (***p<0.001, repeated measures ANOVA with Bonferroni correction). N=4 mice per group. Data represents mean firing rate ± s.e.m.

**(D)** Firing rate of pInsCtx neurons relative to the single auditory pips comprising CS+ $_{fear}$. (n.s., repeated measures ANOVA) N=4 mice per group. Data represents mean firing rate ± s.e.m.

**(E)** Conditioned responses to CS+$_{app.}$. Normalized licking, pupil size, locomotion relative to the start of CS+$_{app.}$. Traces are normalized by the activity 2s before the CS presentation (**p<0.01, student t-test). Sucrose is delivered 3s after the CS+ starts and is indicated by the second blue line. Data represents mean variable ± s.e.m. N=4 mice per group.

**(F)** Conditioned responses to CS-$_{app.}$. Normalized pupil size and locomotion relative to the start of CS-$_{app.}$. Traces are normalized by the activity 2s before the CS presentation (n.s., student t-test). Data represents mean variable ± s.e.m. N=4 mice per group.

**(G)** Conditioned responses to CS+$_{fear}$. Normalized pupil size and locomotion relative to the start of CS+$_{fear}$ (**p<0.01, student t-test). Traces are normalized by the activity 2s before the CS presentation. Data represents mean variable ± s.e.m. N=4 mice per group.

**(H)** Heatmap of the firing rates of heartbeat tuned units in control mice after CS+ $_{app.}$ in control (left) and metoprolol treated mice (right), ordered by peak firing location in the cardiac cycle. (bottom) histogram count of the number of heartbeats tuned units during the cardiac cycle. N=4 mice per group.

**(I)** Same as in (H) but during the CS- $_{app.}$ in control (left) and metoprolol treated mice (right). N=4 mice per group.

**(J)** Same as in (H) but during the CS+ $_{fear}$ in control (left) and metoprolol treated mice (right). N=4 mice per group.
